# Accuracy optimized neural networks do not effectively model optic flow tuning in brain area MSTd

**DOI:** 10.1101/2024.01.26.577393

**Authors:** Oliver W. Layton, Scott T. Steinmetz

## Abstract

Accuracy-optimized convolutional neural networks (CNNs) have emerged as highly effective models at predicting neural responses in brain areas along the primate ventral stream, but it is largely unknown whether they effectively model neurons in the complementary primate dorsal stream. We explored how well CNNs model the optic flow tuning properties of neurons in dorsal area MSTd and we compared our results with the Non-Negative Matrix Factorization (NNMF) model proposed by Beyeler, Dutt, & Krichmar (2016), which successfully models many tuning properties of MSTd neurons. To better understand the role of computational properties in the NNMF model that give rise to MSTd-like optic flow tuning, we created additional CNN model variants that implement key NNMF constraints — non-negative weights and sparse coding of optic flow. While the CNNs and NNMF models both accurately estimate the observer’s self-motion from purely translational or rotational optic flow, NNMF and the CNNs with nonnegative weights yield substantially less accurate estimates than the other CNNs when tested on more complex optic flow that combines observer translation and rotation. Despite their poor accuracy, however, neurons in the networks with the nonnegativity constraint give rise to tuning properties that align more closely with those observed in primate MSTd. Interestingly, the addition of the sparsity constraint has a negligible effect on the accuracy of self-motion estimates and model tuning properties. Across all models, we consistently observe the 90-degree offset in the preferred translation and rotation directions found in MSTd neurons, which suggests that this property could emerge through a range of potential computational mechanisms. This work offers a step towards a deeper understanding of the computational properties and constraints that describe optic flow tuning primate area MSTd.

**Significance Statement:** One of the most exciting developments in visual neuroscience over the past decade is that convolutional artificial neural networks optimized to accurately categorize natural images effectively model neural activity in ventral visual areas of the primate brain. We explored whether accuracy-optimized neural networks account for well-established properties of MSTd, a brain area in the complementary primate dorsal stream that is involved in self-motion perception during navigation. Our findings indicate that such networks depart substantially from MSTd-like tuning, which suggests the computational goal of MSTd may not be to accurately estimate self-motion. We found that adding computational constraints inspired by an existing MSTd model that performs dimensionality reduction on afferent motion signals improves the correspondence with MSTd.

## Introduction

Since the introduction of AlexNet (Krizhevsky, Sutskever, & Hinton, 2012), convolutional neural networks (CNNs) have revolutionized the field of computer vision and have reached the point where they rival or exceed human performance on certain image recognition tasks (Phillips et al., 2018; K. Lee, Zung, Li, Jain, & Seung, 2017; Cireşan, Meier, & Schmidhuber, 2012). Despite their recent impact on computer vision, CNNs have longstanding roots in visual neuroscience (Serre, 2019; Lindsay, 2021). Hubel and Wiesel proposed that the visual system is organized hierarchically based on their seminal discovery of simple and complex cells in cat visual cortex (Hubel & Wiesel, 1962; Hubel, 1982). Fukushima (1980) instantiated this theory as one of the first CNNs called Neocognitron. Remarkably, Neocognitron contains mechanisms that CNNs still use today, including the rectified linear unit activation function (ReLU) that models the nonnegativity and nonlinearity of neuronal firing rates and the max pooling operation that facilitates invariance in pattern recognition.

Given that CNNs contain biologically-inspired mechanisms, it is fascinating that these artificial neural networks have emerged as highly effective at predicting neural responses in areas along the primate ventral stream that include V1 (Cadena et al., 2019; Burg et al., 2021), V4 (Yamins, Hong, Cadieu, & DiCarlo, 2013; Guclu & Van Gerven, 2015), and IT (Yamins & Dicarlo, 2016; Khaligh-Razavi & Kriegeskorte, 2014). CNNs that explain most of the variance of neural responses in ventral stream areas generally achieve the highest image classifcation accuracy on the ImageNet dataset (Yamins et al., 2014; Schrimpf et al., 2018), a collection of more than one million natural images (Deng et al., 2009).

While such accuracy-optimized CNNs have had success in modeling neurons along the ventral stream, it is largely unknown whether they effectively model neurons in the complementary primate dorsal stream. Here we investigated the extent to which CNNs account for well-established tuning properties in dorsal stream area MSTd where neurons demonstrate selectivity to the expansive motion patterns experienced during self-motion (optic flow) (Duffy & Wurtz, 1995; Takahashi et al., 2007). MSTd neurons exhibit systematic tuning to combinations of translational (T) and rotational (R) optic flow (Graziano, Andersen, & Snowden, 1994), which, to first-order approximation, capture any optic flow pattern (Figure 1). It is believed that optic flow sensitivity in MSTd stems from the feedforward integration of local direction and speed signals from MT and other afferent visual areas (Maunsell & Van Essen, 1983; Perrone, 1992; Born & Bradley, 2005).

**Figure 1:**
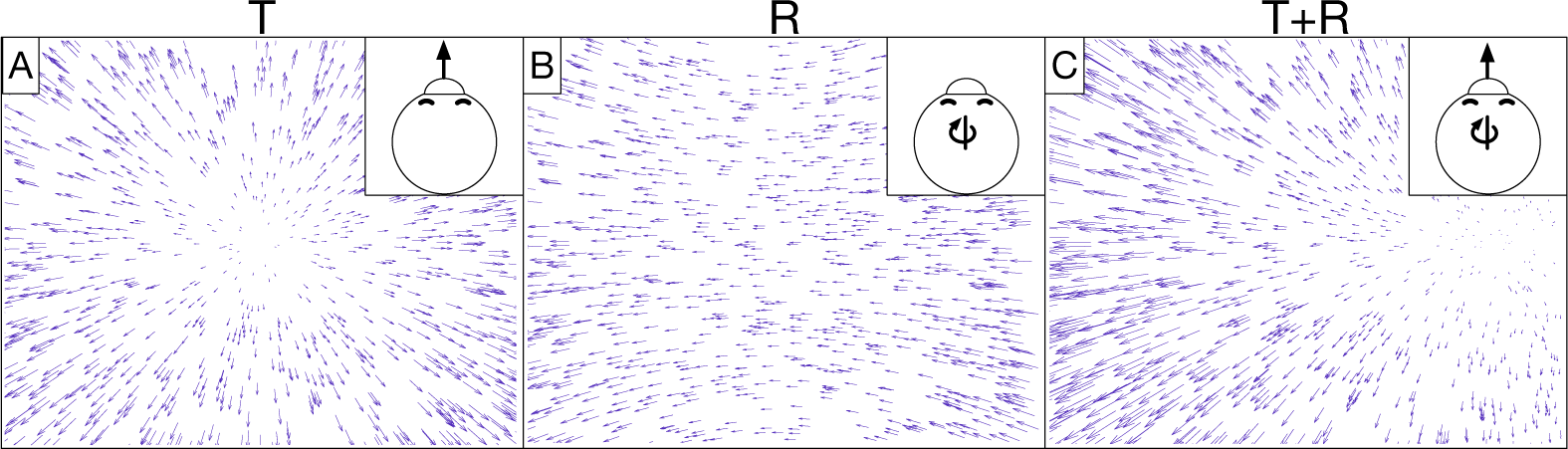
Example optic flow fields generated from simulated self-motion through a 3D dot cloud environment. Self-motion along a straight path of travel creates translational optic flow (T) with motion that radiates from the direction of movement. Eye or head movements yield rotational optic flow (R). (A) depicts translational optic flow from straight-forward self-motion. (B) depicts rotational optic flow corresponding to a rightward eye movement (yaw rotation). (C) depicts the optic flow created from a combination of translation and rotation (T+R). The optic field shown in (C) is the sum of those in (A) and (B).

**Figure 2:**
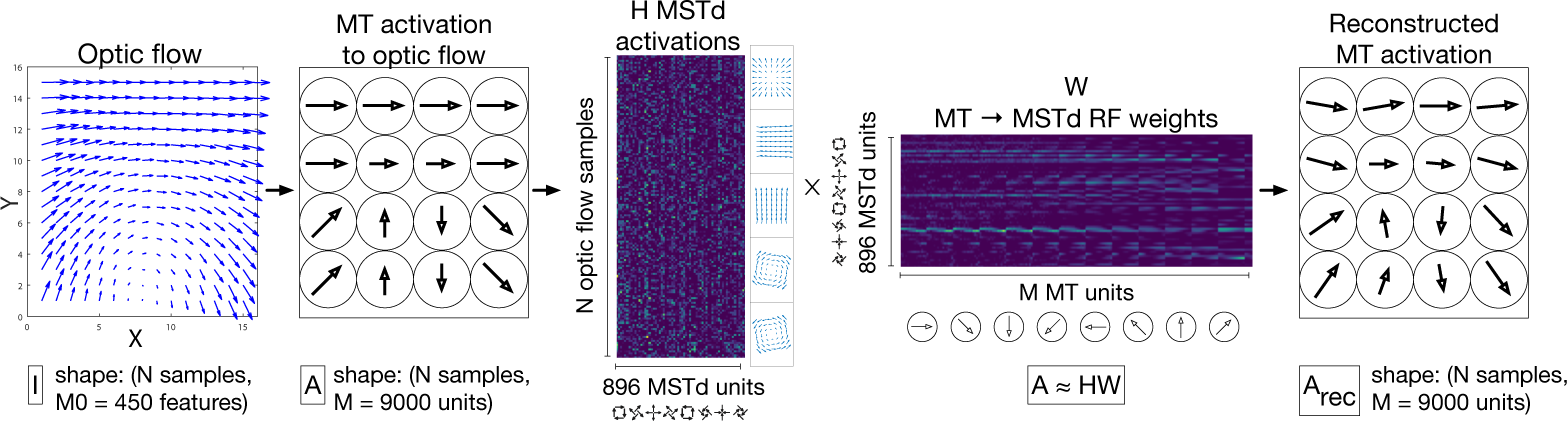
Schematic depiction the NNMF model of MSTd proposed by Beyeler et al. (2016). A population of M = 9000 speed and direction tuned MT units integrate each input 15×15 optic flow field, generating a N × 9000 matrix of activations (A), where N denotes the number of optic flow samples (left two panels). This matrix serves as the input to the NNMF algorithm, which factors A into the matrices H and W (center panels). H corresponds to the nonnegative basis coefficients, which is interpreted as the activations of the 896 model MSTd units to the N optic flow samples. W corresponds to 896 basis vectors, which is interpreted as the MSTd RF connection weights from MT units. The product A*_rec_* = HW is a reconstruction of the MT input activations A (right panel).

In the present study, we optimized CNNs to estimate the translational and rotational self-motion from each optic flow field in a large dataset. We compared the tuning characteristics of neurons in these CNNs to those in the Non-Negative Matrix Factorization (NNMF) model of MSTd proposed by Beyeler and colleagues that reproduces many well-established MSTd properties (Beyeler, Dutt, & Krichmar, 2016). NNMF refers to the process of approximating a matrix A with the product of two other matrices H and W, where all three matrices have nonnegative entries. This forms a basis with which the original matrix may be reconstructed, and, similar to principal component analysis (PCA), the basis has fewer dimensions than the original matrix. Beyeler et al. (2016) apply NNMF to the local motion responses of model MT units to optic flow (A) and interpret the resulting W matrix as MT-MSTd receptive field (RF) connection weights (basis vectors) and H matrix as MSTd unit activations (basis coefficients). This NNMF model can be thought of as a two-layer neural network that, once fit, represents a simple linear model of MT activations (Figure 3A). Unlike PCA, NNMF yields a sparse “parts-based” representation of optic flow patterns wherein many fitted coefficients equal zero (D. Lee & Seung, 1999; Beyeler, Rounds, Carlson, Dutt, & Krichmar, 2019).

**Figure 3:**
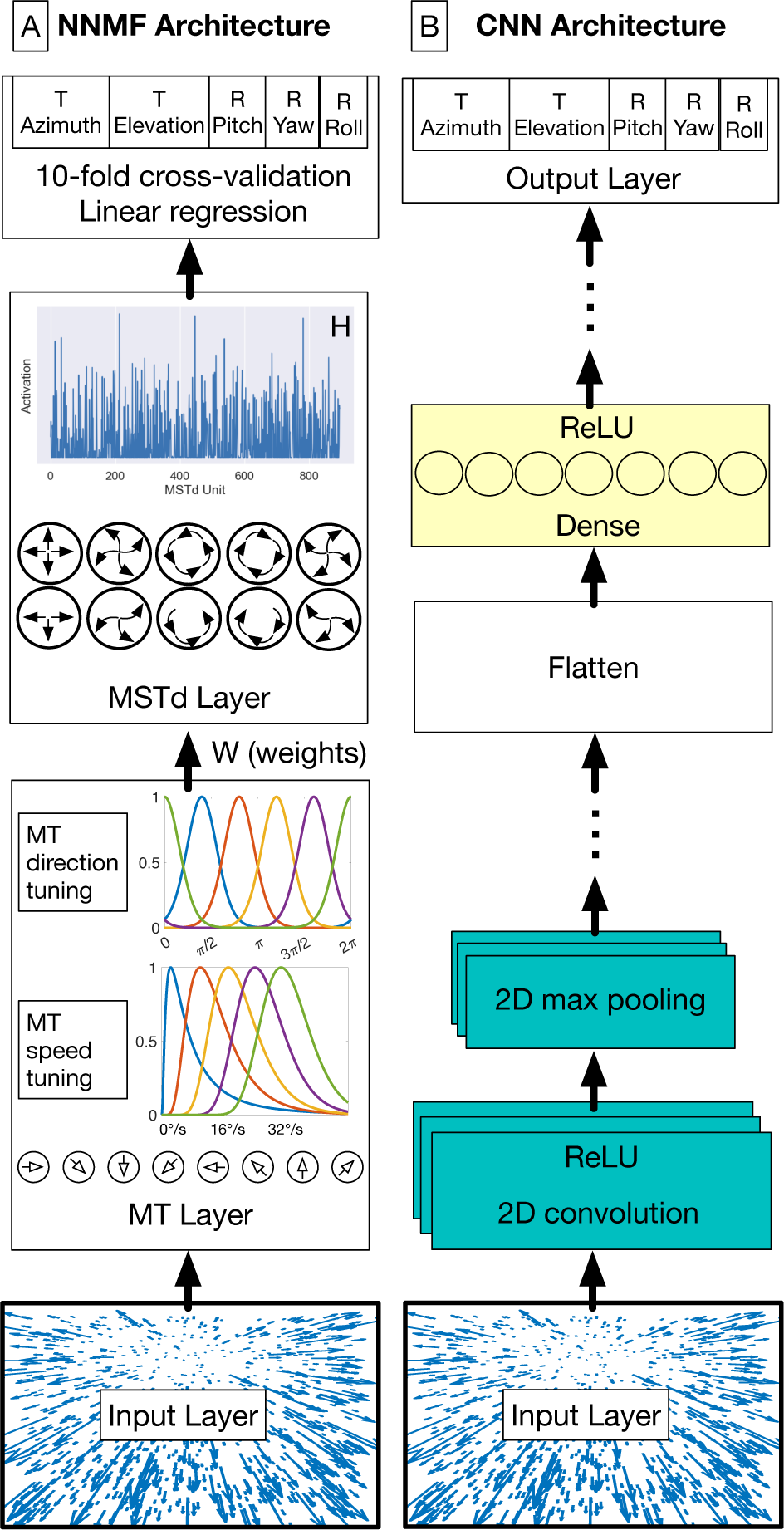
NNMF (A) and CNN (B) model diagrams. (A) In the NNMF model, MT units that are preferentially tuned to a unique combination among 8 directions and 5 speeds integrate optic flow within their RF. MSTd activations emerge by multiplying the MT activations with the MT–MSTd weights obtained from the NNMF algorithm. (B) The first stages of the CNN consist of convolutional units that filter the optic flow field within the RF. This is followed by the max pooling operation, which downsamples the spatial resolution of the optic flow signal. Convolution and max pooling layers may be stacked multiple times before the units are flattened into a 1D representation. These units are connected with one or more layers of densely connected units. The output layer consists of five neurons, one for each self-motion property that the network estimates (the azimuth and elevation of observer translation as well as the pitch, yaw, and roll components of observer rotation). The network minimizes the mean-squared-error (MSE) loss for regression. The MLP model excludes the convolution and max pooling stages (teal).

To better understand what makes NNMF a successful model of MSTd, we also considered CNN variants that incorporate one or both key computational properties of NNMF: nonnegative weights and sparse coding.

## Materials and Methods

We begin by describing the optic flow datasets used to train and evaluate the computational models. We subsequently present the model specifications and end the section with a description of our analyses. To facilitate comparisons with the NNMF model of MSTd, many of our datasets and analyses are based on those of Beyeler et al. (2016) which replicate a number of analyses performed in previous primate studies within a computational modeling framework.

### Optic flow datasets

Our datasets consist of the optic flow generated during simulated self-motion through a scene of 2000 randomly positioned dots. Each data sample corresponds to a single vector field that represents the optic flow produced during an instant of time when the observer moves with 3D translational velocity 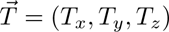 and rotational velocity 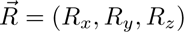. We generate each sample by randomly positioning the dots between 0.1 and 33 m in depth within the 90° field of view of the observer whose eye height is 1.61 m. We project each dot in the world onto the 2D virtual retina using a pin-hole camera model with focal length f = 1.74 cm (Raudies & Neumann, 2013). The projection maps the dot with position (X, Y, Z) within the 3D scene to the point (x, y) on the retina. The following equation specifies the instantaneous optic flow 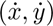 at position (x, y) on the model retina (Longuet-Higgins & Prazdny, 1980):

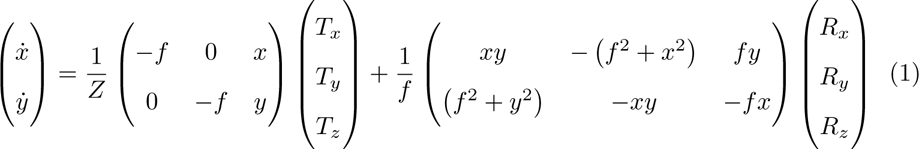

Following Beyeler et al. (2016), we discretized each optic flow sample to a 15×15 vector field. The shape of each optic flow dataset is (N, 15, 15, 2), where N indicates the number of data samples and 2 corresponds to the 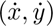 optic flow components.

Table 1 summarizes the datasets used in the present study. Each one adheres closely to the specifications provided by Beyeler et al. (2016). We fit each of the models with the TR360 training set and the remaining datasets serve as test sets to evaluate performance on optic flow that is not used to fit the models (i.e. to test generalization). To create the reported total number of samples, unless noted otherwise, we crossed the independent variables with each other then generated repetitions of these conditions until the number of samples matched the total number. For example, to generate the 6030 TR360 samples for the frontoparallel scene, we crossed T speed (3), R speed (3), and frontoparallel plane depth (5) to obtain 45 samples. We subsequently repeated this process 134 times to obtain the 6030 total fronotoparallel samples. We drew values for indicated random variables (T and R directions in this case) anew when generating each sample so that the dataset does not contain duplicate samples.

**Table 1:**
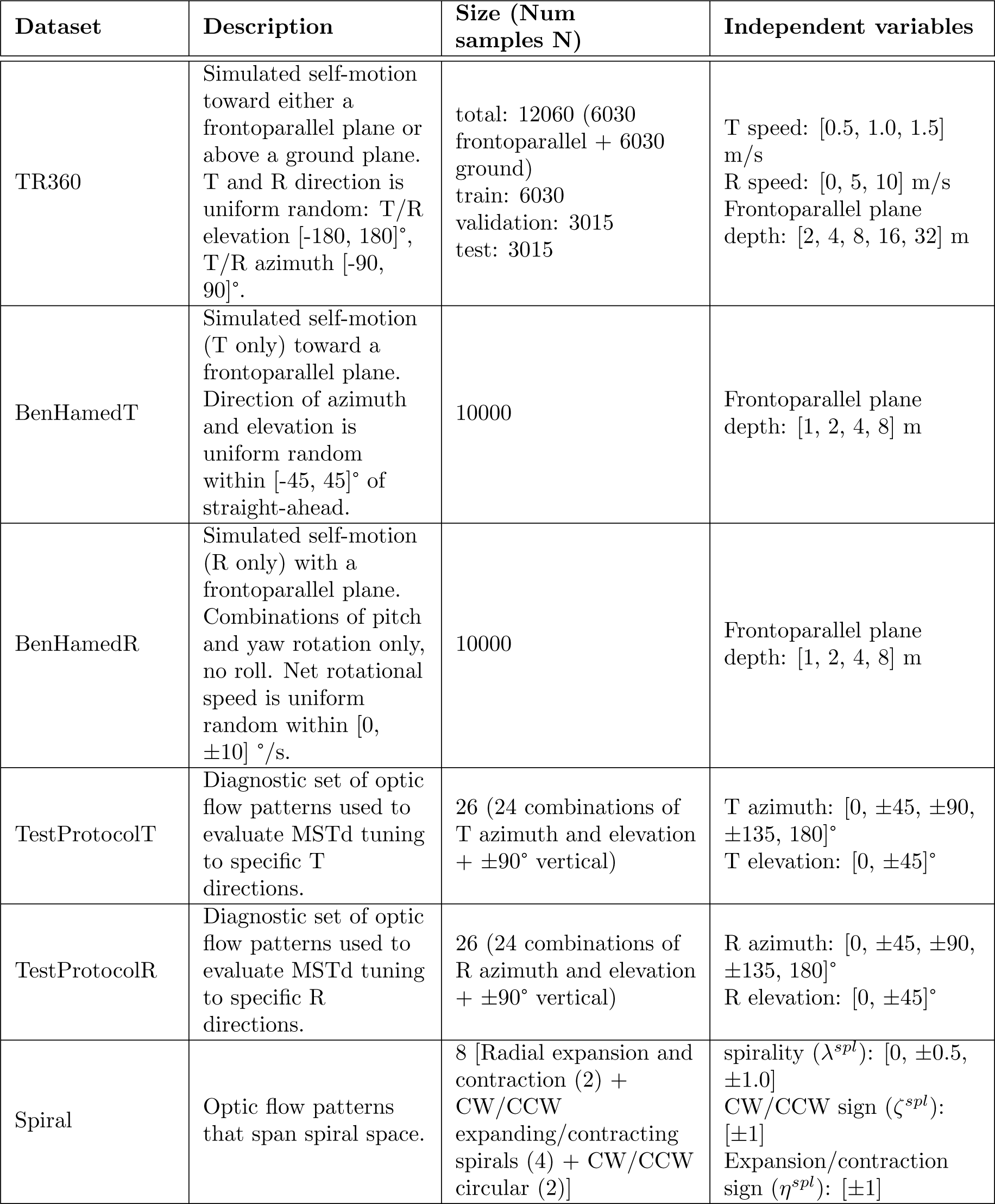
Optic flow dataset specifications. T and R denote translation and rotation, respectively. T indicates optic flow that contains only translation, T+R indicates optic flow with both translational and rotational components. Training set is used to fit network weights, while test set is used to evaluate performance on novel patterns. Uniform indicates sampling from the uniform random distribution with the specified endpoints. Azimuth and elevation angles of 0° correspond to straight-ahead heading. See text for details

| Dataset | Description | Size (Num samples N) | Independent variables |
| --- | --- | --- | --- |
| TR360 | Simulated self-motion toward either a frontoparallel plane or above a ground plane. T and R direction is uniform random: T/R elevation $[-180, 180]^\circ$, T/R azimuth $[-90, 90]^\circ$. | total: 12060 (6030 frontoparallel + 6030 ground)<br>train: 6030<br>validation: 3015<br>test: 3015 | T speed: $[0.5, 1.0, 1.5]$ m/s<br>R speed: $[0, 5, 10]$ m/s<br>Frontoparallel plane depth: $[2, 4, 8, 16, 32]$ m |
| BenHamedT | Simulated self-motion (T only) toward a frontoparallel plane. Direction of azimuth and elevation is uniform random within $[-45, 45]^\circ$ of straight-ahead. | 10000 | Frontoparallel plane depth: $[1, 2, 4, 8]$ m |
| BenHamedR | Simulated self-motion (R only) with a frontoparallel plane. Combinations of pitch and yaw rotation only, no roll. Net rotational speed is uniform random within $[0, \pm 10]^\circ/\text{s}$. | 10000 | Frontoparallel plane depth: $[1, 2, 4, 8]$ m |
| TestProtocolT | Diagnostic set of optic flow patterns used to evaluate MSTd tuning to specific T directions. | 26 (24 combinations of T azimuth and elevation + $\pm 90^\circ$ vertical) | T azimuth: $[0, \pm 45, \pm 90, \pm 135, 180]^\circ$<br>T elevation: $[0, \pm 45]^\circ$ |
| TestProtocolR | Diagnostic set of optic flow patterns used to evaluate MSTd tuning to specific R directions. | 26 (24 combinations of R azimuth and elevation + $\pm 90^\circ$ vertical) | R azimuth: $[0, \pm 45, \pm 90, \pm 135, 180]^\circ$<br>R elevation: $[0, \pm 45]^\circ$ |
| Spiral | Optic flow patterns that span spiral space. | 8 [Radial expansion and contraction (2) + CW/CCW expanding/contracting spirals (4) + CW/CCW circular (2)] | spirality ($\lambda^{spl}$): $[0, \pm 0.5, \pm 1.0]$<br>CW/CCW sign ($\zeta^{spl}$): $[\pm 1]$<br>Expansion/contraction sign ($\eta^{spl}$): $[\pm 1]$ |

In the TR360 dataset, half of samples correspond to simulated self-motion toward a dot-defined frontoparallel plane and the other half correspond to simulated self-motion over a dot-defined ground plane. Following Beyeler et al. (2016), we offset the observer gaze downward by 30° from the horizon in the case of the the ground plane scene (i.e. observer is looking at the ground). We shuffled the order of samples before fitting the models and creating the train/validation/test set splits.

### Computational models

#### Convolutional neural networks

We implemented the CNNs using TensorFlow 2.11 in Python 3.10 (Abadi et al., 2016). We trained the models on a Microsoft Windows 10 workstation equipped with an NVIDIA GeForce RTX 4090 graphics processing unit (GPU) that has 24 GB of RAM. As depicted in Figure 3B, the general CNN architecture processes the optic flow at the input layer with shape (N, 15, 15, 2). The CNN applies 2D spatial convolution to the input signal with stride 1 and zero padded with ‘same’ boundary conditions. This is followed by the ReLU activation function and the 2D max pooling operation. The 2D convolution and max pooling layers are stacked together one or more times. The units in the final max pooling are flattened and connected to one or more layers with dense connectivity and the ReLU activation function. The final stage is a multi-output regression layer with 5 neurons, each representing the self-motion variables that we train the network to estimate: translational azimuth and elevation, as well as rotational pitch, yaw, and roll. Prior to training, we normalized each label separately so that each spans the range [-0.5, 0.5]. The goal of the training process is to minimize the loss incurred in jointly estimating the 5 self-motion variables using the Adam optimizer, configured with default hyperparameters except as noted below. We use mean squared error (MSE) loss for translational elevation (unscaled range: [-90, 90]°) and the rotation variables (unscaled range: [-10, 10]°/s). For the translational azimuth, the network minimizes the following loss function since the range is circular [-180, 180]°:

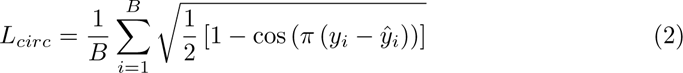

In Eq. 2 B is the mini-batch size, y*_i_* is the translational azimuth label for sample i on the normalized scale, and 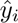 is the normalized predicted translational azimuth value on the normalized scale. This loss function yields the maximum penalty of 1 when the error is 180° offset from the label (e.g. predicting straight-ahead when optic flow of moving straight-back) and 0 penalty when the error is zero or 360° offset from the label. That is, this loss function does not penalize learning angles that deviate by multiples of 360°.

We considered the extent to which several CNN variants modeled MSTd tuning properties:

1. MLP. Multi-Layer Perceptron that possessed the same architecture as the CNN shown in Figure 3B, but without the convolution and max pooling layers (without the teal boxes).
2. CNN L1. The CNN with L1 regularization added to the loss and weights in each layer. The regularization strength was controlled independently in each layer and the values were determined through the optimization process described below.
3. CNN ++. The CNN optimized subject to the constraint that the weights, except for those in the first hidden layer and the output layer, must be non-negative.
4. CNN L1++. The CNN optimized with both L1 regularization and the non-negative weight constraint.

We optimized the neural network architecture and hyperparameters through a two-stage random search process. During the first stage, we conducted a hyperparameter and architecture search for the CNN network based on the ranges shown in Table 2. We fit the CNN to the TR360 training set and recorded the hyperparameters that yielded the smallest validation loss summed across the 5 output neurons. Since our goal was to identify the CNN that best minimizes error when estimating the self-motion variables, we restarted the search with increased upper limits whenever the optimized CNN hyperparameter values encroached on existing limits. During the second stage, we ran independent searches to optimize each the variant networks. We modified the upper limit on the number of possible convolutional layer stacks, dense layers, and units in each layer compared to those listed in Table 2 to match the values used in the optimized CNN from the first stage. This ensured that other optimized models would be no more complex than the baseline CNN to facilitate comparison. We trained all networks with a mini-batch size of 64 samples and early stopping with a patience of 60 epochs. Table 3 shows the optimized hyperparameters values for each of the networks.

**Table 2:** Ranges used in random search for optimal CNN hyper-parameters. Except for the learning rate, hyperparameters were selected on a per-layer basis on every iteration of the search. *L1 regularization was only included in CNN L1 and CNN L1++ networks. **Learning rate was drawn randomly from this set.

| Hyperparameter | Value range |
| --- | --- |
| Number of convolution and max pooling stacks | [1, 4] |
| Number of dense layers | [1, 9] |
| Number of convolutional filters | [2, 600] |
| Number of dense units | [2, 10000] |
| Convolutional unit filter size | [2, 15] |
| Max pooling window size | [2, 4] |
| Max pooling stride length | [1, 3] |
| L1 regularization strength | [0, 1e-3]* |
| Learning rate | [1e-5, 1e-4, 1e-3, 1e-2]** |

**Table 3:** Optimized network hyperparameters. Entries in lists correspond to the values in each relevant layer of the network. For example, [163, 252] means that there are 163 units in the first convolutional layer and 252 units in the second convolutional layer.

| Hyperparameter | CNN | MLP | CNN_L1 | CNN_++ | CNN_L1++ |
| --- | --- | --- | --- | --- | --- |
| Number of convolution and max pooling stacks | 2 | 0 | 1 | 1 | 1 |
| Number of dense layers | 5 | 4 | 5 | 2 | 2 |
| Number of convolutional filters | [163, 252] | 0 | 36 | 148 | 192 |
| Number of dense units | [6856, 5320, 3331, 7787, 5246] | [476, 3617, 3877, 2448] | [4344, 7044, 727, 886, 4660] | [58, 2244] | [5295, 2291] |
| Convolutional unit filter size | [12, 14] | N/A | 2 | 2 | 2 |
| Max pooling window size | [2, 2] | N/A | 2 | 2 | 2 |
| Max pooling stride length | [2, 1] | N/A | 2 | 3 | 2 |
| L1 regularization strength | 0 | 0 | [1e-6, 1e-7, 1e-6, 1e-14, 1e-11, 1e-11, 1e-13] | 0 | [1e-9, 1e-9, 1e-15, 1e-16] |
| Learning rate | 1e-5 | 1e-3 | 1e-5 | 1e-3 | 1e-3 |

#### Non-negative matrix factorization model

We implemented and configured the NNMF model according to the specifications of Beyeler et al. (2016), except where noted otherwise.

The first stage of the model (Figure 3A) transforms each 15×15 optic flow field into motion signals in model area MT. Model MT neurons exhibit sensitivity to the speed and direction of optic flow signals over time. We simulated 9000 (N*_MT_*) model MT neurons: 8 preferred directions and 5 preferred speeds centered on each of the 15×15 positions in the optic flow input (9000 = 8 × 5 × 15 × 15).

Tuning to the direction θ(x, y) present at position (x, y) within the RF of a neuron relative to the preferred direction θ*_pref_* obeys a von Mises distribution:

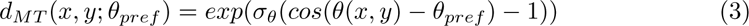

where σ*_θ_* = 3° indicates the bandwidth of the direction tuning, which was set to approximate the ≈ 90° full-width at half-maximum found in MT cells (Britten & Van Wezel, 1998; Beyeler et al., 2016).

The tuning of each model MT neuron to the optic flow speed (ν) at position (x, y) within the RF obeys a log-normal distribution (Nover, Anderson, & Deangelis, 2005):

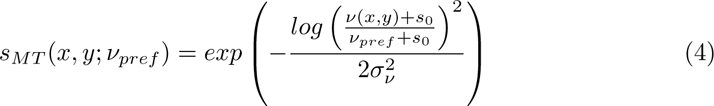

where σ*_ν_* defines the speed tuning bandwidth, s_0_ defines a non-negative offset parameter to prevent the singularity in the logarithm at 0, and ν*_pref_* defines the preferred speed of the model neuron. We set σ*_ν_* = 1.16 and s_0_ = 0.33 to match the median values obtained from neural data Nover et al. (2005). The 5 model MT preferred speeds are 2, 4, 8, 16, and 32°/s (Beyeler et al., 2016; Nover et al., 2005).

The activation of each model MT neuron is the product of the direction and speed inputs within the RF:

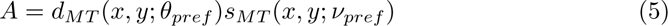

We used NNMF to decompose the MT activation matrix A into the product of two other matrices HW, all of which must have non-negative entries:

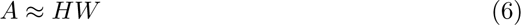

The MT matrix A has shape (N, M), where N corresponds to the number of optic flow samples and M corresponds to the number of MT units. The matrix H has shape (N, K) represents the activations of the K = 896 model MSTd units to each sample. The matrix W has shape (K, M) represents the weights between the M = 9000 MT units and the K = 896 model MSTd units. NNMF minimizes the MSE reconstruction loss L*_nnmf_* between A*_rec_* = HW and A

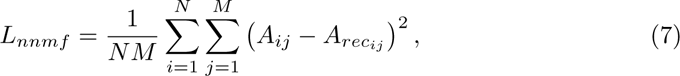

subject to the non-negativity constraint that all entries H*_ik_ ≥* 0 and W*_kj_ ≥* 0. We implemented NNMF in TensorFlow and used gradient descent with the Adam optimizer (learning rate: 1e-3) to minimize L*_nnmf_*. We enforced the non-negativity constraint on each training epoch through the ReLU function.

NNMF has several free hyperparameters that must be specified. First, the matrices H and W require initialization. We used uniform random values between 0 and 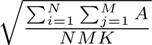, as used in the Scikit-learn library implementation of NNMF (Pedregosa et al., 2011). Second, the number of basis vectors in the decomposition K must be selected. Following Beyeler et al. (2016), we repeatedly fit NNMF 14 times, using 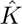 = 64 basis vectors in each instance. This allows the model to incorporate variability in the fitted basis vectors due to the random initiation of H and W. We concatenated the matrices fit from each of the 14 NNMF fits to obtain K = 896. Third, NNMF requires convergence criteria for the iterative optimization of Eq. 7 to be set. This is important since fitting NNMF for too many iterations risks overfitting the training set. We stopped the fit when the absolute difference in the loss obtained between successive training epochs was at most 1e-4. This is the default tolerance value used in the MATLAB function nnmf, which Beyeler et al. (2016) used to fit their model (we are assuming they used default parameter values since they do not state otherwise). We required the NNMF fit to run for at least 2 epochs. Fourth, an algorithm must be selected by which NNMF is fit. We used gradient descent since it allowed us to fit NNMF using the same paradigm as the neural networks and our TensorFlow implementation allowed us to train quickly on the GPU. We obtained similar results using the alternating least-squares algorithm, which is what Beyeler et al. (2016) used.

We used the following equation to compute MSTd activations H*_test_* in the fitted NNMF to N*_test_*novel stimuli.

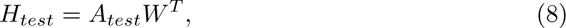

where A*_test_*is the MT activations to the novel stimuli (shape: (N*_test_*, M)) and W *^T^* is the transpose of the MT–MSTd fitted NNMF basis vectors (shape: (M, K)).

### Analyses

We focused our analyses of the units in the final hidden layer of the CNNs and the MSTd layer of the NNMF algorithm (H*_test_*). Unless noted otherwise, we excluded neurons that did not produce nonzero activation to any of the test optic flow patterns (“unresponsive neurons”/“dying ReLUs”) from the analysis (Glorot, Bordes, & Bengio, 2011; Lu, Shin, Su, & Karniadakis, 2019). All our analyses focused on model outputs on the test sets, novel optic flow not used to fit the models.

#### MSTd receptive field maps

We used using the TestProtocolT and TestProtocolR diagnostic datasets to characterize the tuning of model units to different translation and rotation directions. First, we interpolated the unit’s activations obtained to the N = 26 optic flow patterns within each diagnostic dataset on a regular azimuth-elevation mesh. We subsequently evaluated the interpolation at 40 sample points made up of 8 azimuths (0–360° in 45° increments) and 5 elevations (−90–90° in 45° increments). Finally, we created a heat map showing a Lambert cylindrical equal-area projection: the horizontal axis corresponds to the azimuthal angle θ, the vertical axis corresponds to the elevation angle ψ transformed as sin(ψ), and the color corresponds to the unit activation.

In addition to generating heat map plots showing individual unit tuning, we created composite heat maps showing the activation at each sample point averaged across the entire model population.

#### Translation and rotation tuning preferences

We established the translation and rotation tuning preferences of each unit using the TestProtocolT and TestProtocolR datasets (see Table 1). We estimated each neuron’s preferred tuning using the population vector method (Georgopoulos, Schwartz, & Kettner, 1986; Beyeler et al., 2016). This involves summing the product between each unit activation and the corresponding optic flow translation or rotation labels (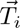 or 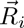), represented as 3D Cartesian unit vectors.

#### Population tuning width

To characterize the T or R tuning width of each model unit, we considered the interpolation obtained on the TestProtocolT and TestProtocolR datasets. We evaluated the interpolation at 100 azimuths and 50 elevations (3.6° increments along either axis). We defined a unit’s tuning width as the Euclidean distance between where unit achieves its maximum and half-maximum activation.

#### Peak heading discriminability

To quantify how well individual model units discriminate between similar headings, we presented each model with 24 optic flow patterns with equally spaced translation directions spanning 0–360° (step size: 15°) in the horizontal plane (0° elevation). We fit separate cubic splines with 1000 equally spaced sample points to the set of activations produced by each model unit. We evaluated the first derivative of the spline to measure the discriminability at different reference directions of translation. Following Gu, Watkins, Angelaki, and Deangelis (2006) and Beyeler et al. (2016), we report the peak heading discriminability, defined as the heading direction at which the spline derivative reaches its maximum. To mitigate edge and circularity effects, we padded the endpoints with activations produced to additional optic flow stimuli with 20 translation direction steps reaching ±300° on either side of the 0–360° central interval.

#### Heading and rotation tuning index

The translation direction (heading) that generates the strongest response represents the heading preference for a particular model unit. The heading tuning index (HTI) measures the selectivity of each unit’s heading tuning. A neuron with a strong heading preference (HTI ≈ 1) activates only to a narrow range of heading directions, while a neuron that activates to a broad range of headings exhibits a weak preference (HTI ≈ 0).

We computed the HTI for each model unit j on the TestProtocolT dataset according to (Beyeler et al., 2016; Gu et al., 2006):

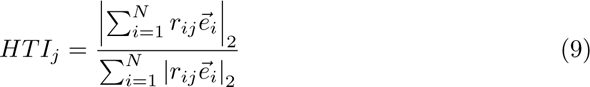

where r*_ij_* denotes the activation of the j^th^ model unit in response to the i^th^ stimulus, 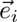 denotes the heading direction for the i^th^ stimulus as a 3D Cartesian unit vector, was the Cartesian heading direction of the ith stimulus in unit vector form, and | · |_2_ denotes the L^2^ Euclidean norm. This equation produces HTI values ranging from 0 (no directional tuning) to 1 (strong directional tuning).

Moreover, we used using Eq. 9 to compute a tuning index that measures each neuron’s rotation selectivity that we refer to as the rotation tuning index (RTI). We computed the RTI of each neuron based on the activations obtained on the rotation-only TestProtocolR diagnostic dataset, substituting the 3D Cartesian rotation unit vector (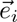) for sample i in Eq. 9.

#### Sparseness metrics

We assessed the sparseness (s) of model MT unit activity (r) using the following metric (Vinje & Gallant, 2000):

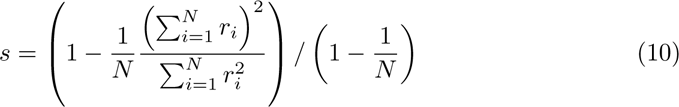

where s is a metric ranging from 0 to 1, with 0 indicating a dense code where every neuron is always active and 1 indicating a local code where a single neuron is active for each stimulus. A sparse coding represents in the middle ground, where a subset of neurons activate to any given stimulus. We use this equation to calculate two measures of sparseness. The first is population sparseness, the proportion of model units active for a single stimulus, where r*_i_* represents the activity of the i^th^ model unit to a stimulus and N is the total number of model units. The second is lifetime sparseness, the proportion of stimuli a single model unit is active for, where r*_i_* represents the activity of a model unit to the i^th^ stimulus and N is the total number of stimuli. The population and lifetime sparseness metrics reflect average values for all stimuli and model units, respectively.

### Software Accessibility

We implemented and simulated NNMF and the CNNs in Python using the NumPy (Harris et al., 2020), SciPy (Virtanen et al., 2020), Pandas (McKinney, 2010), Seaborn (Waskom, 2021), and TensorFlow (Abadi et al., 2016) libraries. The model code will be released on GitHub.

## Results

Inspired by the success of accuracy-optimized CNNs at modeling neural activity in primate ventral stream areas, we examined the extent to which CNNs capture optic flow tuning in dorsal stream area MSTd. We optimized CNNs to estimate the observer’s visual translation and rotation on a 6030 sample optic flow dataset (TR360) composed of 3D linear translation sampled from all possible directions and 3D rotation sampled from a range of commonly encountered speeds (0–10°/s) (see Table 1). In addition to a “baseline” CNN model (see Materials and Methods), we simulated CNN variants that implement two key characteristics of the NNMF model of Beyeler and colleagues whose properties closely emulate many well-established characteristics of MSTd: sparse coding of optic flow and non-negativity in connectivity weights that gate afferent signals. Our aim was to examine how these computational principles influence the correspondence with MSTd properties. The three CNN variants that we considered are:

1. CNN with L1 regularization (LASSO) in each layer (Tibshirani, 1996), which promotes spareness in each neuron’s weights (CNN L1).
2. CNN optimized with non-negative weight constraints imposed on each internal network layer (CNN ++).
3. CNN with both L1 regularization and non-negative weight constraints (CNN L1++).

To better understand the potential importance of the convolution operation on model properties, we compared the CNNs to a multi-layer perception (MLP), which possesses the same architecture except that it lacks convolution and max pooling layers early in the network.

We begin by presenting the accuracy with which CNNs perform this task before comparing CNN translation and rotation tuning properties with those of primate area MSTd.

### Accuracy of self-motion estimates

We assessed the accuracy with which the neural networks estimate the 3D translation and rotation of the observer on a 3015 sample test set, novel optic flow stimuli never encountered during training. The CNN estimates translation and rotation with a high degree of accuracy — the CNN achieves mean absolute errors (MAEs) of 7.1° (Figure 4B) and 1.3° (Figure 4D), respectively. The mean squared error (MSE) in estimating the corresponding self-motion parameters are 322.9°^2^ (Figure 4A) and 2.5°^2^ (Figure 4C). As Figure 4A–D indicates, the accuracy achieved by the MLP is comparable to that of the CNN. While CNN L1 performs similarly to both the CNN and MLP, networks with the non-negative weights estimate translation and rotation with far less accuracy (CNN ++, CNN L1++; Figure 4A–D). Inspection of the individual predictions reveals qualitative differences: whereas CNN, MLP, and CNN L1 yield a high concentration of estimates near true values along the unity line (Figure 4G), predictions from the CNN ++, CNN L1++ are more dispersed and deteriorate markedly when estimating backward translation (≈±180°). Combining L1 regularization and the non-negative weight constraint does not improve accuracy compared to CNN ++ (Figure 4I).

**Figure 4:**
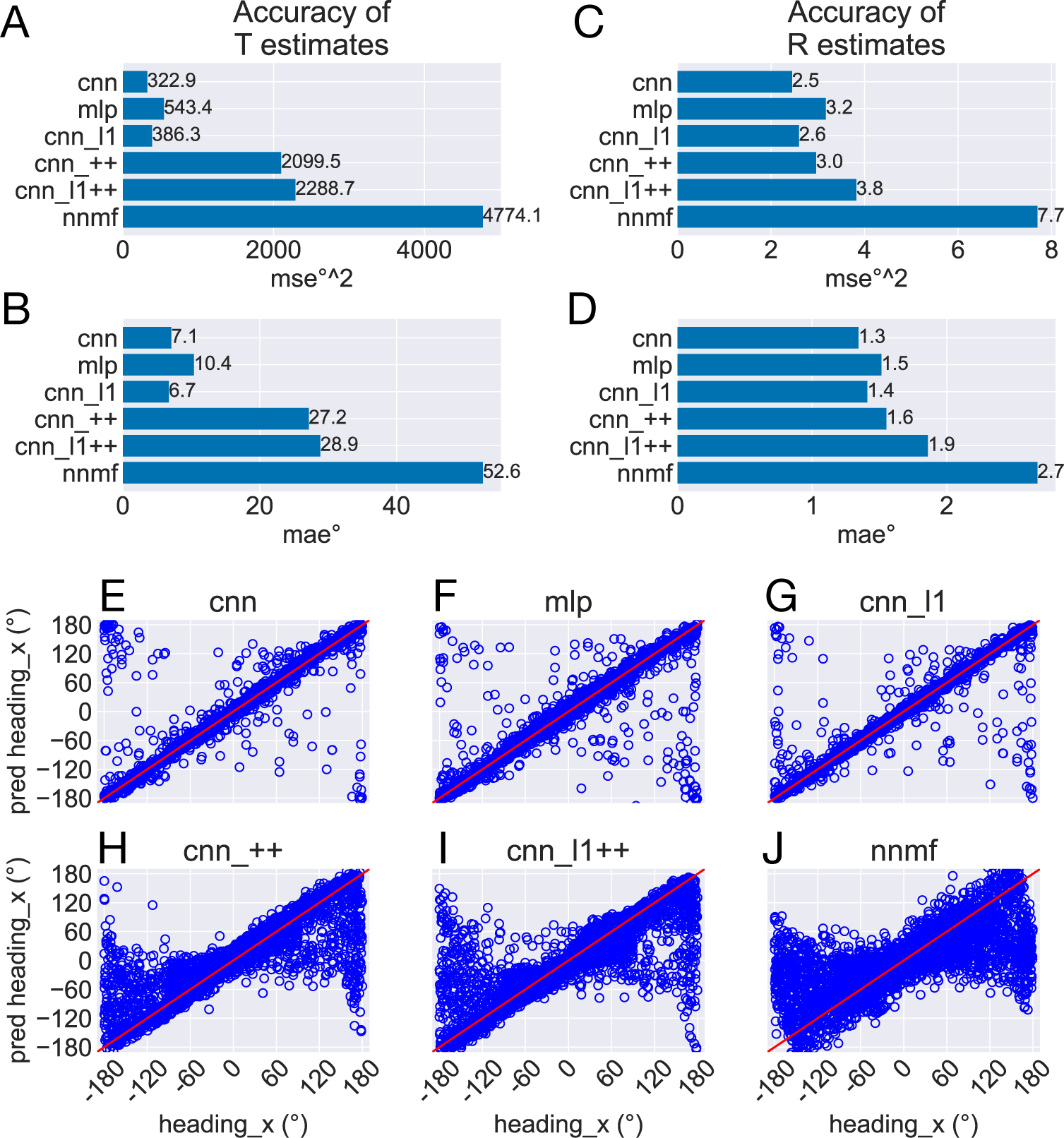
Test accuracy on the TR360 optic flow dataset obtained by the CNN and NNMF models. MSE represents mean squared error and MAE represents mean absolute error. (A–B) Accuracy of translational self-motion estimates achieved by each model. (C–D) Accuracy of rotational self-motion estimates achieved by each model. (E–J) Scatter plots focus on the model predictions for individual optic flow test samples in the case of azimuthal translation (“heading x”). The x axis corresponds to the true value, the y axis corresponds to the value predicted by the model, and the red curve shows the unity line (no error). Note that headings of 180° and −180° correspond with the same direction (backward).

For comparison, we simulated the original NNMF model of Beyeler and colleagues (Beyeler et al., 2016). To do this, we fit a linear regression to the NNMF model MSTd activations obtained on the training set stimuli, and, similar to the CNNs, we assessed the accuracy of estimates obtained on the test set. Figure 4A–D shows that the NNMF model yields substantial translation and rotation errors, with approximately twice the MSE and MAE obtained by the CNNs that yield the least accurate estimates (CNN ++, CNN L1++). Figure 4J reveals that the individual NNMF predictions more closely resemble those produced by CNN ++ and CNN L1++ than those without the non-negativity constraint. It is noteworthy that NNMF garnered poor accuracy when estimating the TR360 test but not the training set (MAE ± STD, 13.97°, ± 18.04). Thus, generalization is poor in the NNMF model.

The poor accuracy of self-motion estimates decoded from NNMF is somewhat surprising given that Beyeler et al. (2016) report only ≈ 5° and 1° mean error when decoding translation and rotation, respectively, from the optic flow stimuli of Ben Hamed, Page, Duffy, and Pouget (2003). These are novel stimuli not used to fit the model. The discrepancy in accuracy could stem from the fact that the Ben Hamed stimuli were considerably simpler than the TR360 dataset simulated here. The Ben Hamed stimuli consist of two separate datasets: either pure translation within ±45° of straight-ahead (BenHamedT dataset) or pure rotation (BenHamedR dataset). This contrasts with the TR360 dataset that contains combined translation and rotation along any 3D direction. Without refitting the models (i.e. the weights reflect learning on the more general TR360 dataset), we decoded self-motion from the CNNs and the NNMF model on the Ben Hamed datasets to determine whether task difficulty could account for the large difference in accuracy. Following Beyeler et al. (2016), we decoded from the NNMF model by fitting a separate linear regression for each self-motion label and assessing the mean validation accuracy through a 10-fold cross validation procedure. We carried out this process for the BenHamedT and BenHamedR datasets independently. To facilitate comparison with the CNNs that were trained end-to-end on the TR360 dataset with a supervised learning paradigm, we lesioned the weights between the last hidden layer and the output layer and fit separate linear regressions to the last hidden layer activations through the same 10-fold cross validation procedure that was used with the NNMF model.

Figure 5 shows the MAE obtained by cross-validation when the models estimate the horizontal (x) and vertical (y) translational (heading) components in the BenHamedT dataset (Figure 5A) and the corresponding rotational components in the BenHamedR dataset (Figure 5B). With the exception of the MLP in the case of y heading, both NNMF and the neural networks estimated self-motion with a high degree of accuracy — MAEs of several degrees or less. The substantial gap in accuracy between the CNN variants with and without the non-negativity constraint (CNN ++ and CNN L1++) obtained on the TR360 dataset (Figure 4) largely vanishes on the Ben Hamed datasets. It is important to consider that the BenHamedT dataset contains headings ranged from −45° to 45° in both the horizontal and vertical directions and BenHamedR dataset contains rotational speeds that range between −10 and 10°/s. Together, these findings suggest that the NNMF model encodes optic flow accurately when decoding pure translation and rotation over limited extents, but incurs substantial error when estimating self-motion from more complex combinations of translation and rotation. The fact that most models yield highly accurate estimates on Ben Hamed datasets supports the notion that task difficulty explains the discrepancy between NNMF accuracy on the TR360 and Ben Hamed datasets.

**Figure 5:**
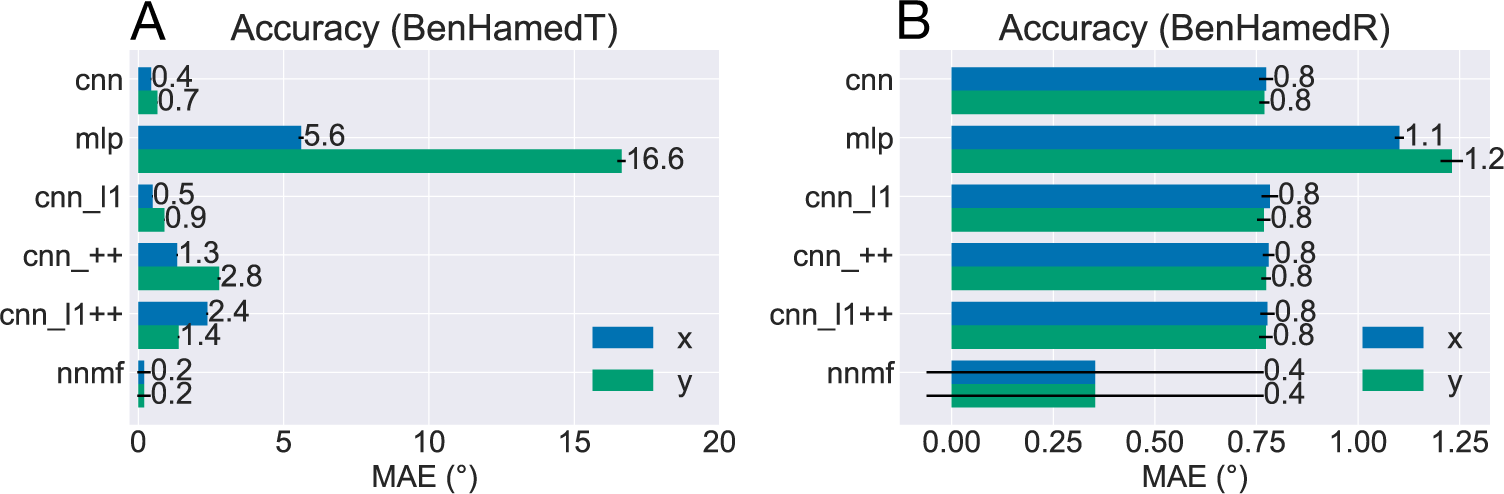
Accuracy of each model on the BenHamedT and BenHamedR datasets. The BenHamedT dataset consists of optic flow corresponding to translation-only self-motion and the direction of translation varies −45–45° in both x and y. The BenHamedR dataset consists of optic flow corresponding to rotation-only self-motion with −10–10°/s speeds. Error bars show 95% confidence intervals (CIs) on the MAE estimated by 10-fold cross-validation.

To establish whether the disparity in accuracy achieved by NNMF across the TR360 and Ben Hamed dataset stems from its non-negative and sparse “parts-based” representation, we applied the PCA algorithm to the MT optic flow activations and performed the same decoding procedure. PCA is another dimensionality reduction algorithm that does not enforce non-negativity and does not yield a parts-based encoding. We found that PCA configured with 64 basis vectors (i.e. same as NNMF) yields the same pattern of results as NNMF — 5142°^2^ and 9°^2^/s MSEs when estimating translation and rotation, respectively, on TR360 test samples as well as 0.4° and 0.5°/s MAEs in both x and y when estimating translation and rotation, respectively, on the Ben Hamed stimuli. We conclude that the poor NNMF generalization on the TR360 dataset is not necessarily the consequence of a non-negative and sparse encoding.

### Translation tuning profiles

Now we turn our analysis to characterizing the tuning of model units to translational optic flow and drawing comparisons with primate MSTd. The heatmaps in Figure 6 show the translation tuning profile of four units randomly selected from each model alongside the tuning profiles of two MSTd neurons from Takahashi et al. (2007). We generated each heatmap by considering the neural responses of a single model unit to 26 translational optic flow stimuli with regularly spaced azimuths and elevations from the TestProtocolT diagnostic dataset (see Materials and Methods). To facilitate comparison with existing neurophysiological and modeling work, we adopt the same coordinate system as Takahashi et al. (2007) and Beyeler et al. (2016) where 0° and 180° azimuth correspond to leftward and rightward translation, respectively, and −90° and 90° elevation correspond to upward and downward translation, respectively (see bottom-right schematic in Figure 6). In some cases the tuning profiles of single model units share qualitative similarities with the MSTd neuron exemplars (e.g. MSTd Neuron 2, NNMF Neuron 1, CNN Neuron 2).

**Figure 6:**
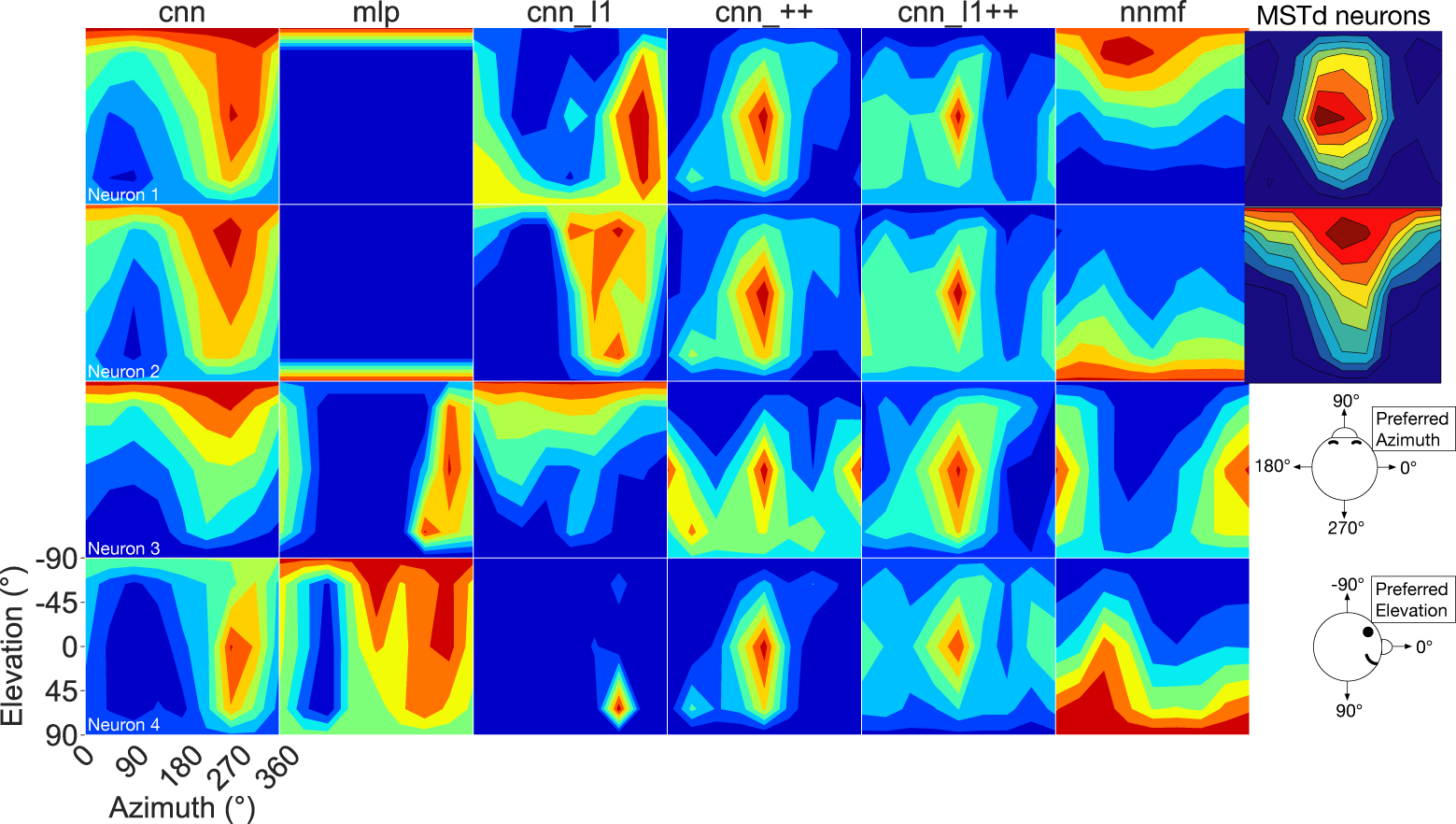
3D translation tuning profiles for four randomly selected neurons (rows) in each model (columns). Translation directions are expressed respect to azimuth and elevation angle (°). Warmer colors correspond to stronger activation to specific combinations of azimuth and elevation angles and cooler colors correspond to weaker activation. The translation tuning profiles for two MSTd neurons from Takahashi et al. (2007) are depicted in the rightmost column. The bottom-left diagram schematizes the translation direction coordinate system.

To characterize translation responses more generally across the population, we summed the activations obtained to each translation direction and generated population translation tuning profiles for each model (Figure 7). The CNN, MLP, and CNN L1 models (Figure 7A–C) responded maximally to 270° azimuths (backward translation) and minimally to 90° azimuths (forward translation). Strikingly, the CNNs with the non-negativity weight constraint inspired by the NNMF model (Figure 7D–E) demonstrate 90° shifted maximal responses across the population — the peak is concentrated at 180° azimuth (leftward translation). The CNN ++ and CNN L1++ model population response peaks resemble that of the NNMF model (Figure 7F). It is noteworthy that the population activations produced by CNN ++, CNN L1++, and MLP models are more tightly concentrated around 0° elevation (translation parallel to ground plane) than the CNN and CNN L1 models, which produced large activations over a wider range of elevation angles. While the NNMF population peaks at 0° elevation like the former subset of models, it also responds to a wide range of elevation angles.

**Figure 7:**
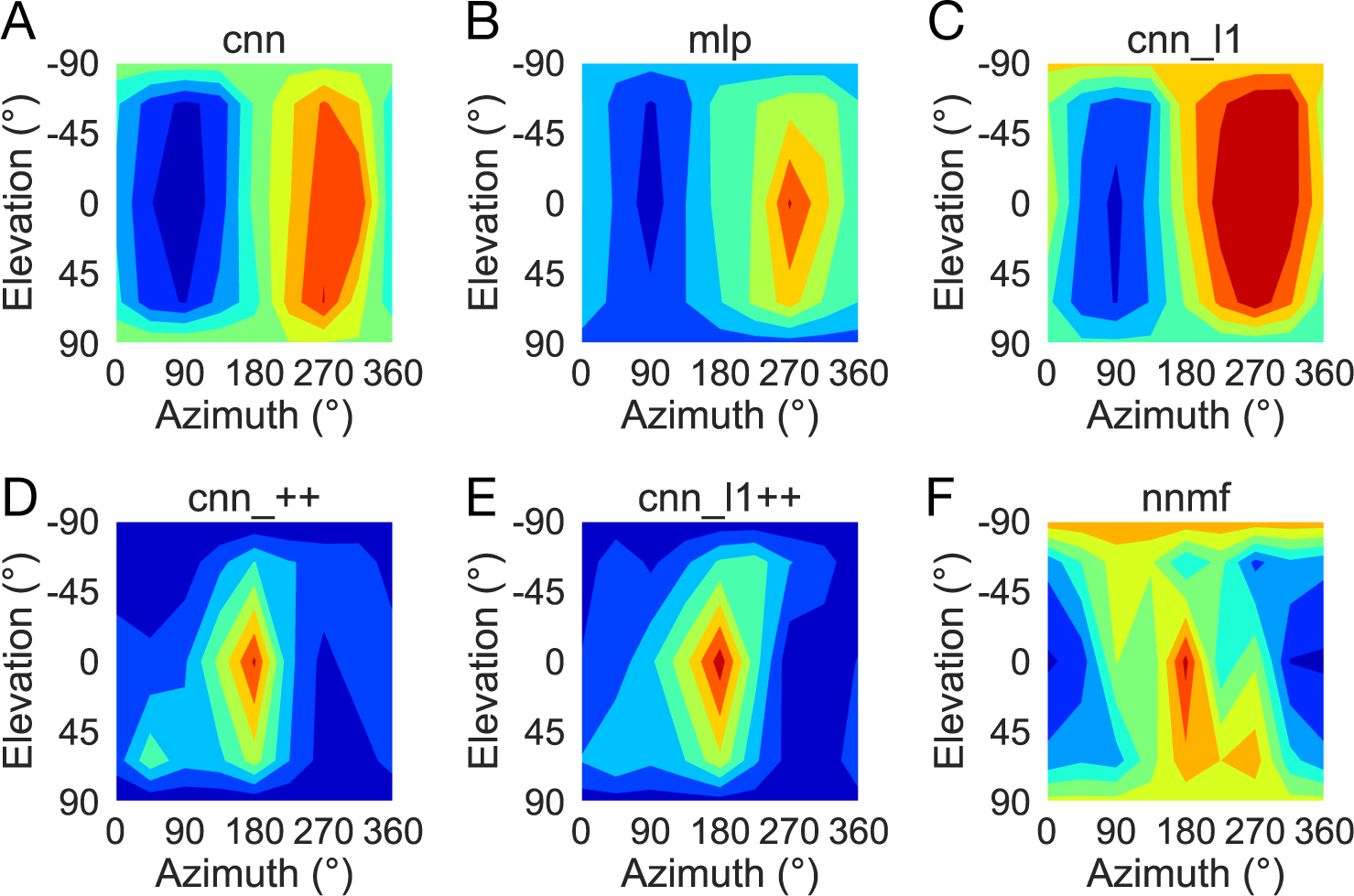
Population translation tuning profiles for each model. Same plotting conventions and coordinate system as Figure 6.

### Translation direction preferences

Following Takahashi et al. (2007) and Beyeler et al. (2016), we characterized the translation tuning preference of each model unit using population vector decoding (see Materials and Methods). Each scatter plot in Figure 8 shows the preferred translational azimuth and elevation angle of single neurons in a particular model. The histograms to the right and above each scatter plot show the marginal distribution of preferred azimuth and elevation angles, respectively, across the population. From the scatter plots and histograms it is clear that the translation preferences of NNMF model units (Figure 8G) exhibit the greatest consistency with those of MSTd neurons (Figure 8H). A key characteristic of MSTd preferences for translational azimuth angle is the relative paucity of neurons that prefer fore-aft self-motion directions (90°, 270°) compared to lateral directions (0°, 180°; Figure 8H). While few units in the CNN, MLP, and CNN L1 models prefer ≈90° azimuth (forward) translation, many units in these models possess ≈270° (backward) azimuthal preferences, a departure from the MSTd data. The large number of neurons with backward translation preferences in these models are consistent with their population tuning profiles (Figure 7A–C). Despite their divergence from the overall distribution of MSTd azimuthal preferences, Table 4 reveals that the CNN (20%), MLP (13%), and CNN L1 (18%) came closest among the simulated models to the percentage of MSTd neurons found by Takahashi et al. (2007) to have translation preferences within 30° of the lateral axis (20%).

**Figure 8:**
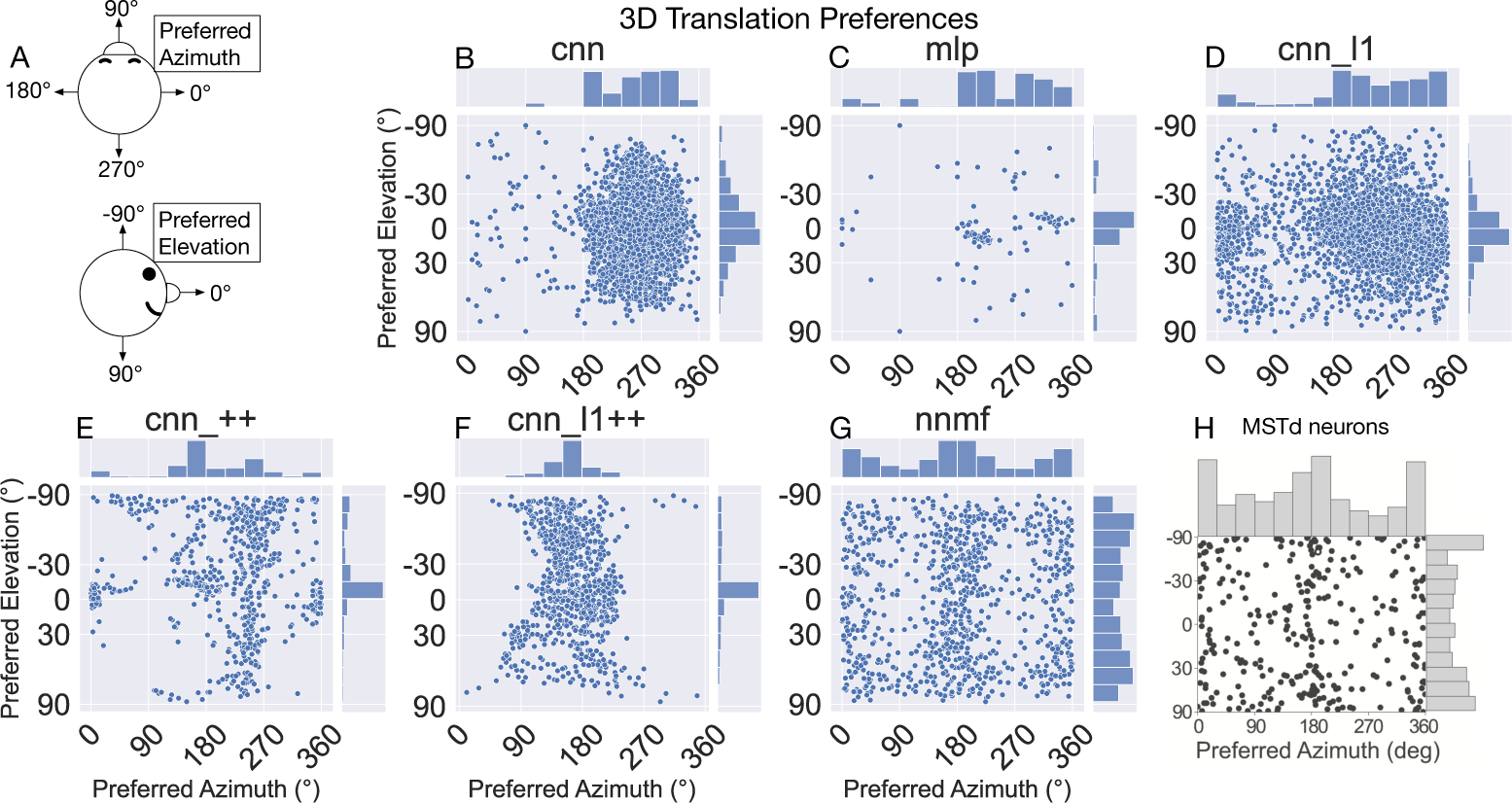
The 3D translation direction preferences of single neurons in each model. The preferred direction of each neuron is expressed as its preferred combination of azimuth and elevation angles. The bar charts on top and to the right of each scatter plot depict histograms (12 bins, bin width: 30°), showing the marginal distribution of neurons with different preferred azimuth and elevation angles, respectively. (A) Schematic depicting the the coordinate system, which is the same as in Figure 6. (H) 3D translation direction preferences of MSTd neurons from Takahashi et al. (2007).

**Table 4:**
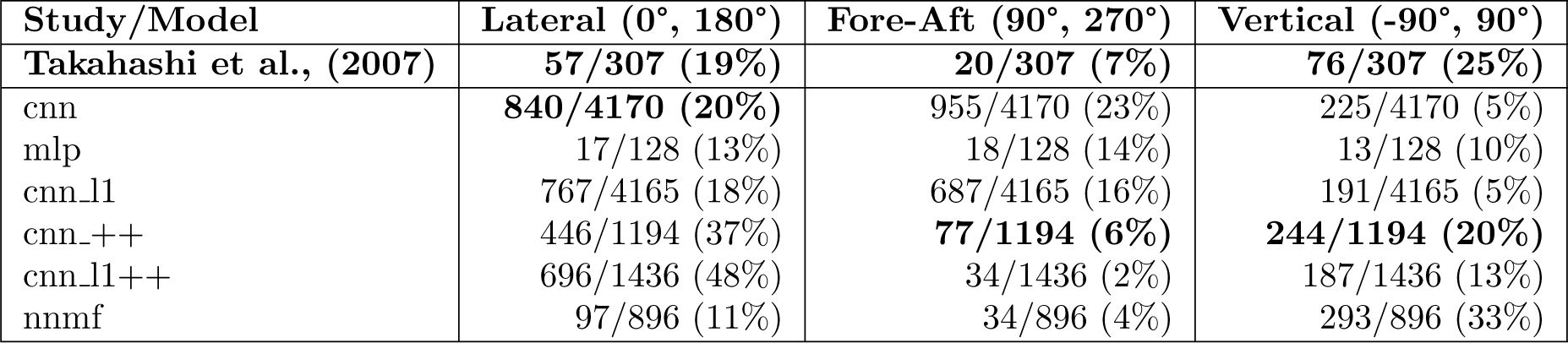
The percentage of neurons in each model with preferred translation directions within 30° of the lateral (leftward-rightward), fore-aft (forward-backward), and vertical (upward-downward) axes. Each bold entry shows the closest percentage to that reported by Takahashi et al., (2007) for MSTd.

The networks with the non-negativity constraint (CNN ++, CNN L1++) exhibit greater consistency with MSTd azimuthal preferences, with a peak in preferences around 180° (leftward) and paucity of preferences in the fore-aft directions. Indeed, the proportion of units that have preferences within 30° of the fore-aft translation axis in the CNN ++ (6%) and CNN L1++ (2%) models comes close to matching the percentage of MSTd neurons (7%). With the exception of CNN L1, none of the neural networks yield the high concentration of azimuthal preferences around 0/360° (rightward) that is present in the MSTd data (Figure 8H).

With respect to elevation, MSTd neurons are most likely to prefer large angles (upward and downward translation; Figure 8H). However, all of the neural network models yield the opposite pattern — most of the neurons prefer ≈0° elevation (Figure 8B–F). Of the CNNs, CNN ++ possesses the highest proportion of neurons with large elevation preferences, albeit mostly for ≈-90°. Indeed, Table 4 shows that CNN ++ comes the closest (20%) out of the simulated models to the percentage of MSTd neurons found by Takahashi et al. (2007) to have translation preferences within 30° of the vertical axis (25%; Table 4).

In summary, while the CNN, MLP, and CNN L1 models have the most MSTd-like percentage of units that prefer nearly lateral translation directions, these models deviate from MSTd preferences due to the predominance of units that prefer backward translation. The CNNs with the non-negativity constraint closely match the percentage of MSTd neurons that prefer translation along the fore-aft direction and produce more MSTd-like azimuthal preferences than the other neural networks. However, substantial discrepancies remain in all the CNNs; the NNMF model best approximates MSTd translation preferences.

### Translation tuning width

The preceding analysis characterizes the translation direction that elicits the maximal response in each unit, but it does not capture the spatial extent of the tuning. Figure 9 plots the tuning width at half maximum for single units in each model (see Materials and Methods). As indicated in Figure 9A, we adopt a different coordinate system to facilitate comparison with MSTd translation tuning width data from Gu, Fetsch, Adeyemo, Deangelis, and Angelaki (2010) (Figure 9H). In this coordinate system, the −90° and 90° correspond to leftward and rightward translation, respectively, and 0° and ±180° correspond to forward and backward translation, respectively. The kernel density estimates on the top of each scatter plot reflect the aforementioned backward bias in CNN, MLP, and CNN L1 translation preferences (±180°) and the preponderance of preferences in the CNN ++ and CNN L1++ models for leftward translation (−90°). All of the neural networks produce the pattern in the MSTd data where neurons that prefer headings within 45° of the straight-ahead generally yield narrower tuning widths (green markers) and those that prefer more peripheral headings (blue markers to the left and right side of the green markers) have greater variability and range in their tuning widths (Figure 9B–F). However, only the NNMF model exhibits the tendency of MSTd neurons for the minimum tuning widths to be at least 45°, regardless of the preferred direction. Interestingly, the subpopulation of neurons that prefer nearly straight-ahead headings (green markers) in the MLP all had exquisitely narrow and similar tuning widths of ≈22.5°.

**Figure 9:**
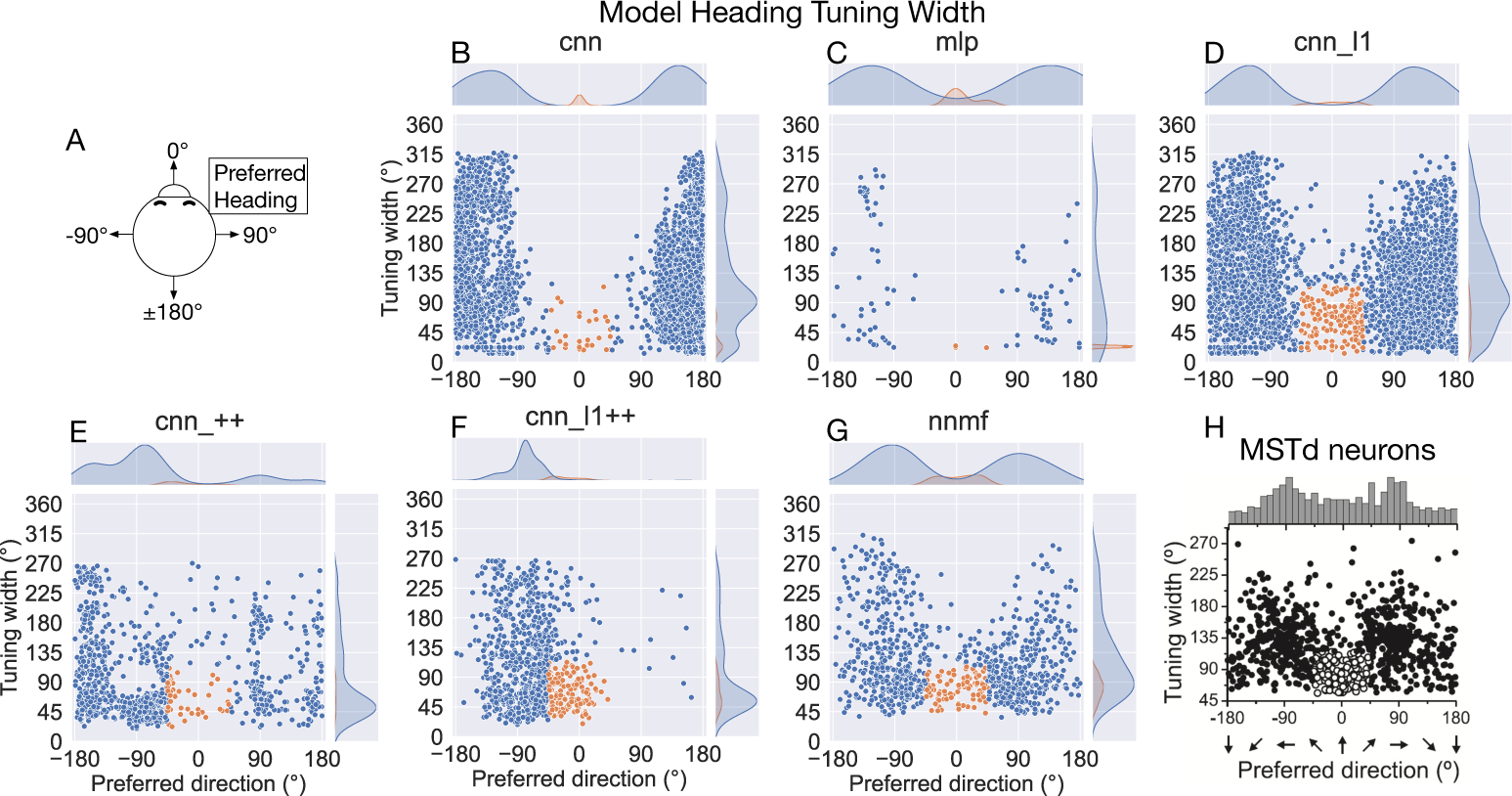
Translation tuning width at half maximum of each model neuron. (A) Schematic of coordinate system, which is different than in Figure 6 to facilitate comparison with MSTd data from Gu et al. (2010) (H). Filled curves on the top and right side represent the marginal kernel density estimates. As depicted in (H), scatter plots show neurons with preferred heading within 45° of straight-ahead and tuning widths < 115° in green.

### Heading discriminability and strength of tuning

While MSTd neurons across the population exhibit lateral (leftward and rightward) bias in their translation preference, they possess peak discriminability for straight-ahead, and to a lesser extent, backwards headings (Gu et al., 2010) (Figure 10G). To assess the extent to which the models reproduce this tendency, we computed the heading at which each model units demonstrates maximal discriminability (see Material and Methods). The CNN, MLP, and CNN L1 models yield qualitatively similar distributions, with peak discriminability at backward headings (≈±180°). While MSTd neurons demonstrate elevated discriminability to backward headings, the peak in discriminability for straight-ahead headings is virtually absent in these three models. Taken together with Figures 7–8, these models largely prefer and are most sensitive to backward headings. Interestingly, the CNNs with the non-negativity constraint both capture the peak discriminability at straight-ahead headings and produce lesser peaks for backward headings, albeit to a weaker extent than in the MSTd population (Figure 10D–E). While the NNMF model yields a peak close to straight-ahead and lesser peaks at greater eccentricities (Figure 10F), its four distinct peaks are punctuated by sharp drops in peak discriminability at the cardinal axes (0°, 90°, ±180°), a pattern that does not appear in the MSTd data (Figure 10G).

**Figure 10:**
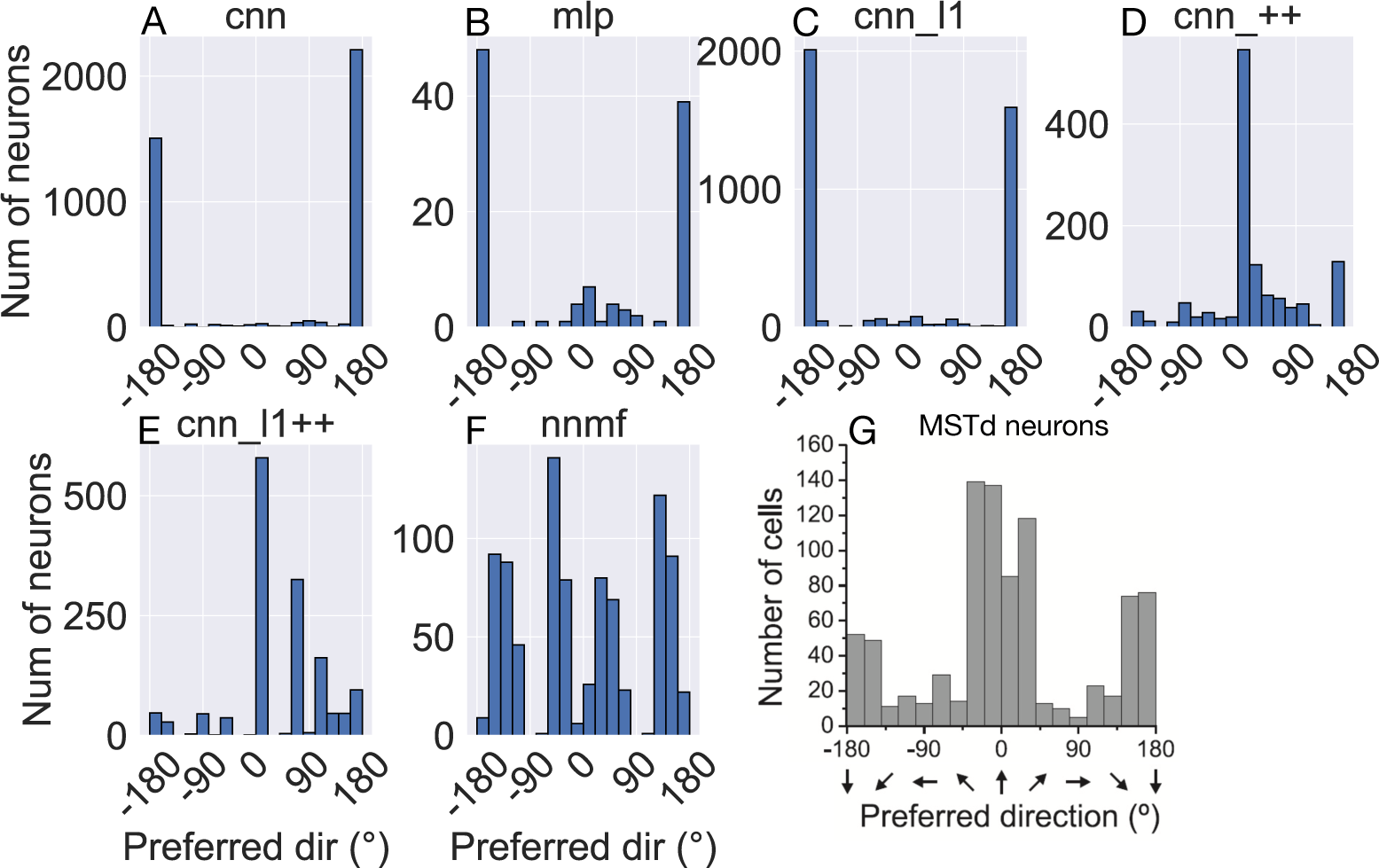
Histograms (18 bins, width: 20°) showing peak heading discriminability of units in model. Optic flow stimuli involve translation in the horizontal plane without variations in elevation or rotation. Coordinate system of Figure 9 is used. (G) Peak heading discriminability of MSTd neurons from Gu et al. (2010).

Figure 11 shows the heading tuning index (HTI) computed on the TestProtocolT diagnostic dataset (see Materials and Methods). The HTI quantifies the strength of a unit’s heading tuning and ranges between 0 and 1 (Gu et al., 2006). A value of 0 indicates weak heading tuning while a value of 1 indicates strong heading tuning (i.e. unit selectively activates to only a single heading). Gu et al. (2006) obtained a mean HTI of 0.48 ± 0.16 SD for their MSTd population. Due to the differences in the HTI distribution shapes, we report median HTIs for each model population. The CNN (median ± SD, 0.42 ± 0.19) and CNN L1 (0.39 ± 0.22) demonstrate the best agreement with the MSTd HTIs. Consistent with their narrow tuning widths (Figure 9C), MLP neurons are more heading selective (95% CI: [0.64, 0.72]) than the CNN [0.42, 0.43] and CNN L1 [0.37, 0.38]. Neural populations in CNN ++ [0.25, 0.27], CNN L1++ [0.16, 0.18], and NNMF [0.26, 0.28] exhibit less overall heading selectivity.

**Figure 11:**
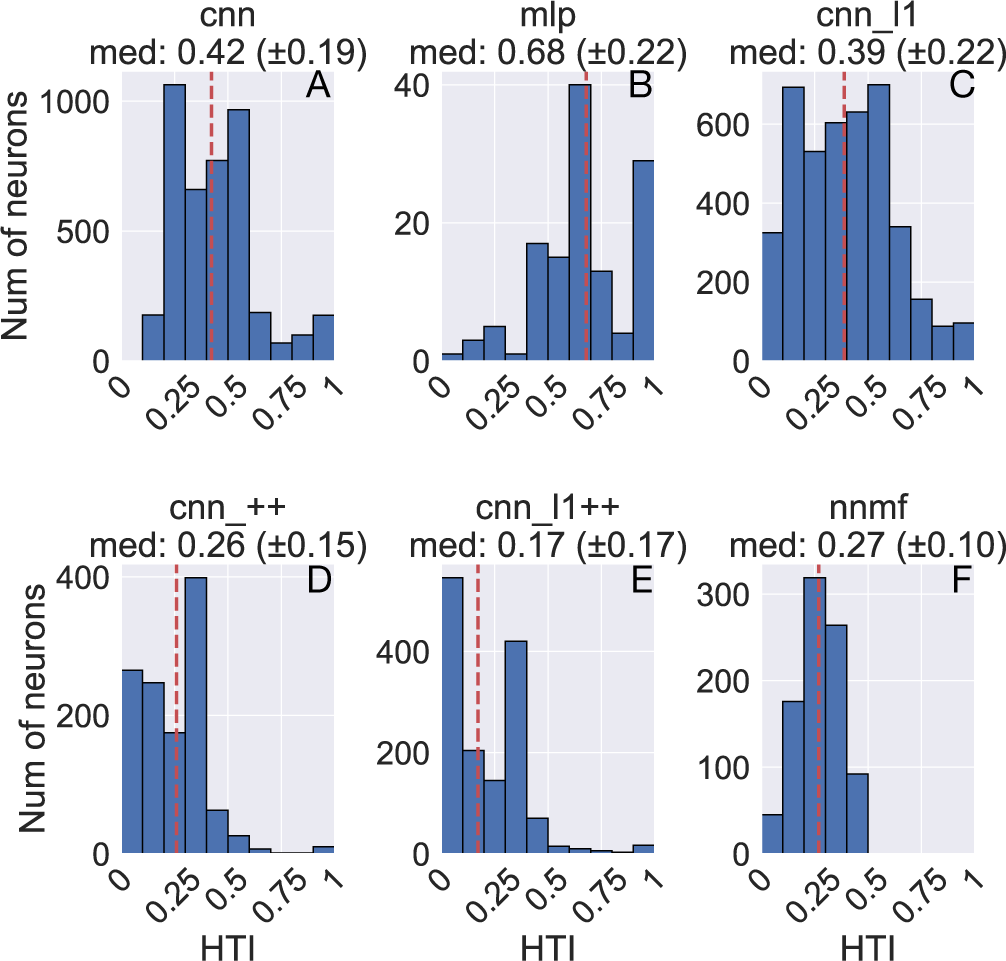
Histograms (10 bins) showing the heading tuning index (HTI) of units in each model. The values above each plot show the median HTI across the population as well as its standard deviation. The dashed vertical line corresponds to the median HTI.

In summary, the correspondence between the models and MSTd heading discriminability and strength of tuning is mixed. While the CNN ++, CNN L1++, and NNMF models capture key aspects of MSTd peak heading discriminability (Figure 10D–F), these models demonstrate weaker heading selectivity than neurons in MSTd (Figure 11D–F). Conversely the CNN and CNN L1 models yield more compatible HTI values (Figure 11A,C), but deviate in the peak heading discriminability (Figure 10A,C).

### Rotation direction preferences

Next we focus on model tuning to rotational optic flow. Figure 12 shows a sample of the diverse rotation tuning profiles derived from the single unit activations to the TestProtocolR diagnostic dataset. Figure 13 shows the preferred directions of rotation for single units in each model population. The coordinate system follows the conventions of Figure 8, except each pair of preferred azimuth and elevation angles now indicate the direction about which visual rotation induces the maximal activation. Consistent with Beyeler et al. (2016) and MSTd rotation preferences (Figure 13H), the NNMF model population peaks at around 0/360° (left) and 180° (right) in azimuth preferences. However, the NNMF distribution of elevation preferences is broad and does not peak around vertical axes of rotation (±90°). Peaks in CNN, MLP, and

**Figure 12:**
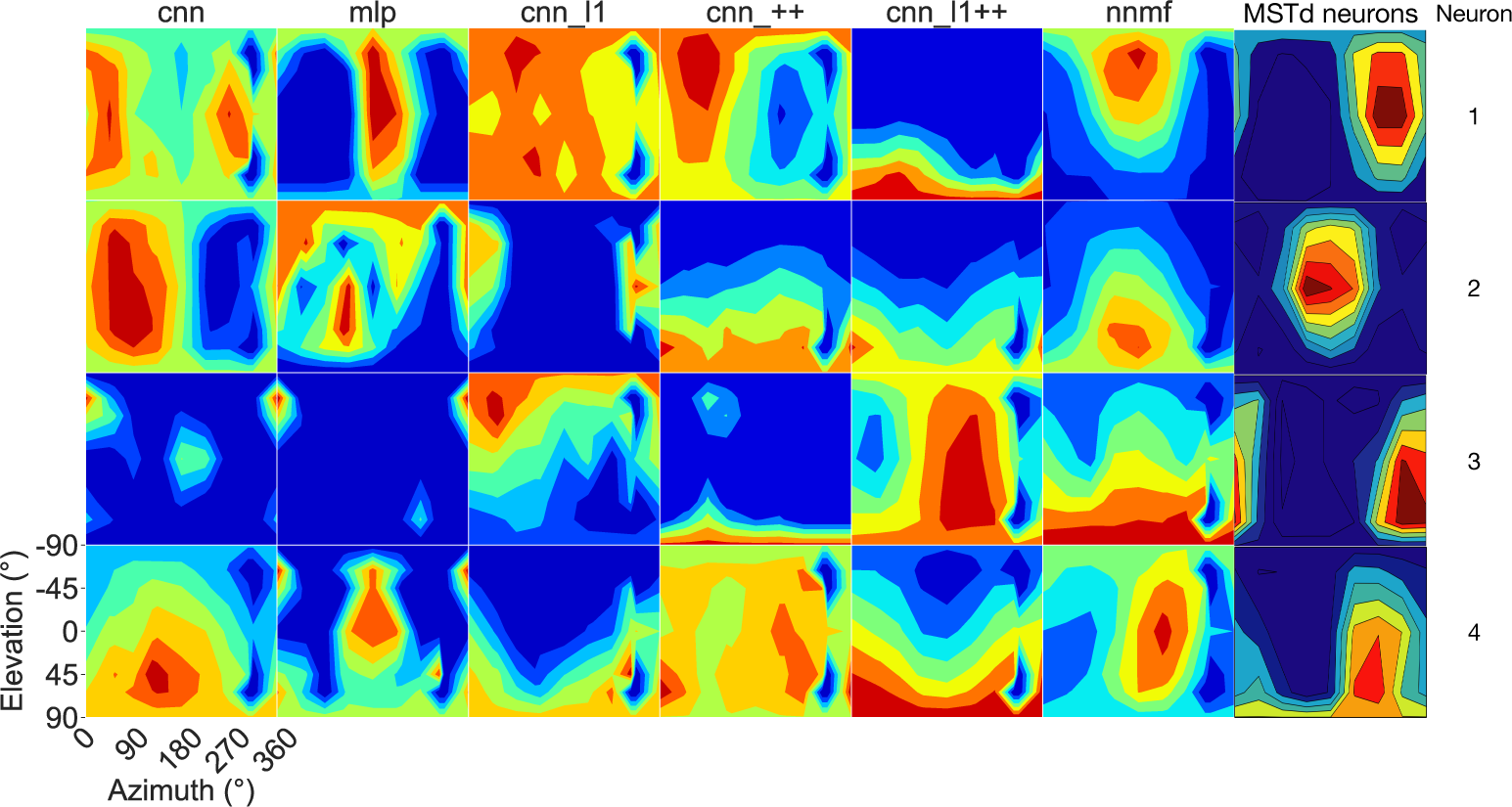
3D rotation tuning profiles for four randomly selected neurons (rows) in each model (columns). Same format and coordinate system as Figure 6. The rotation tuning profiles for four MSTd neurons from Takahashi et al. (2007) are depicted in the rightmost column.

**Figure 13:**
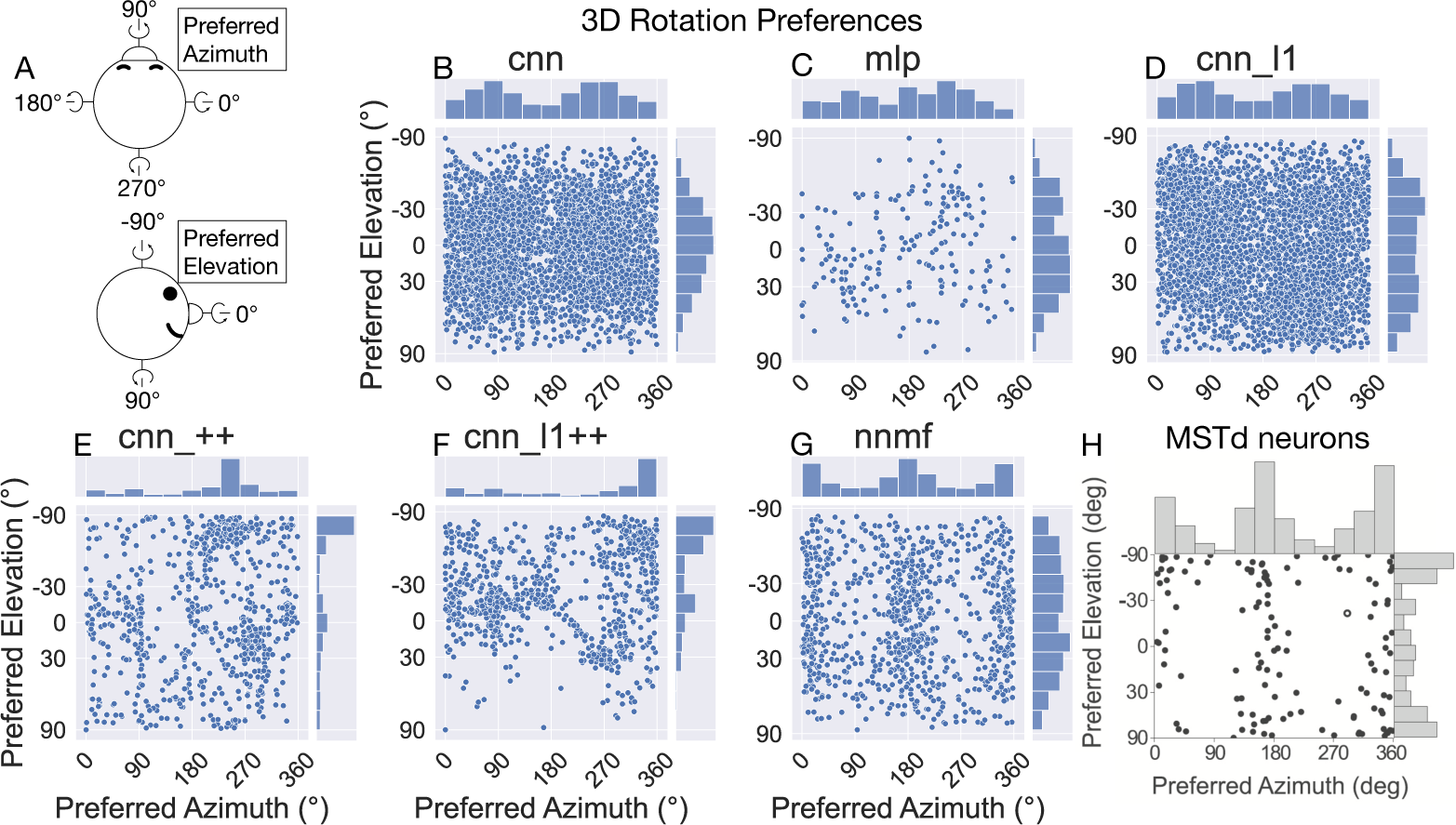
The 3D rotation direction preferences of single neurons in each model. Same format as Figure 8. (A) Schematic depicting the the coordinate system. (H) 3D rotation direction preferences of MSTd neurons from Takahashi et al. (2007).

CNN L1 population azimuth preferences (Figure 13B–D, see marginal histograms) appear shifted by ≈90° compared to the MSTd population (Figure 13H). Similar to the NNMF model population, the distribution of preferred rotation elevation angles is broad and not peaked at ±90°. Unlike the translation case (Figure 8), the non-negativity constraint did not consistently improve the agreement between the CNNs and the MSTd data. While the CNN L1++ model and the MSTd data have consistent peaks in preferred azimuth and elevation, the model distribution deviates in its unimodal shape and the preferred azimuths in the CNN ++ appear shifted by 90° like in the CNN, MLP, and CNN L1 models. Overall, rotation preferences in the NNMF model produce the best agreement with those of MSTd neurons. In support of this, the NNMF model offers the best match to MSTd in the percentage of neurons that have rotation preferences within the 30° of the vertical yaw axis (NNMF: 19% vs. MSTd: 28%), the horizontal pitch axis (15% vs. 21%), and the depth roll axis (4% vs. 1%; see Table 5).

**Table 5:** The percentage of neurons in each model with preferred rotation directions within 30° of the vertical (yaw), lateral (pitch), and depth (roll) axes. Each bold entry shows the closest percentage to that reported by Takahashi et al., (2007) for MSTd.

| Study | Yaw | Pitch | Roll |
| --- | --- | --- | --- |
| **Takahashi et al., (2007)** | **36/127 (28%)** | **27/127 (21%)** | **1/127 (1%)** |
| cnn | 276/4216 (7%) | 229/4216 (5%) | 1273/4216 (30%) |
| mlp | 15/223 (4%) | 15/223 (7%) | 32/223 (14%) |
| cnn_l1 | 634/4128 (15%) | 193/4128 (5%) | 739/4128 (18%) |
| cnn_++ | 643/1159 (56%) | 46/1159 (4%) | 155/1159 (13%) |
| cnn_l1++ | 684/1405 (49%) | 58/1405 (4%) | 197/1405 (14%) |
| nnmf | **166/896 (19%)** | **130/896 (15%)** | **33/896 (4%)** |

Figure 14 shows the difference in each model unit’s preferred translation and rotation angle. Interestingly, all the models yield median differences of ≈90°, consistent with MSTd neurons from Takahashi et al. (2007) (Figure 14G). That is, the shift in preference occurs despite the considerable differences in computational mechanisms that distinguish the models. It is noteworthy that the models with the non-negativity constraint (CNN ++, CNN L1++, NNMF) yield sharp, more distinct peaks at ≈90° (Figure 14C–F) more akin to the MSTd distribution than the other neural networks (Figure 14A–C). The NNMF model best captures the relatively narrow spread of the MSTd distribution.

**Figure 14:**
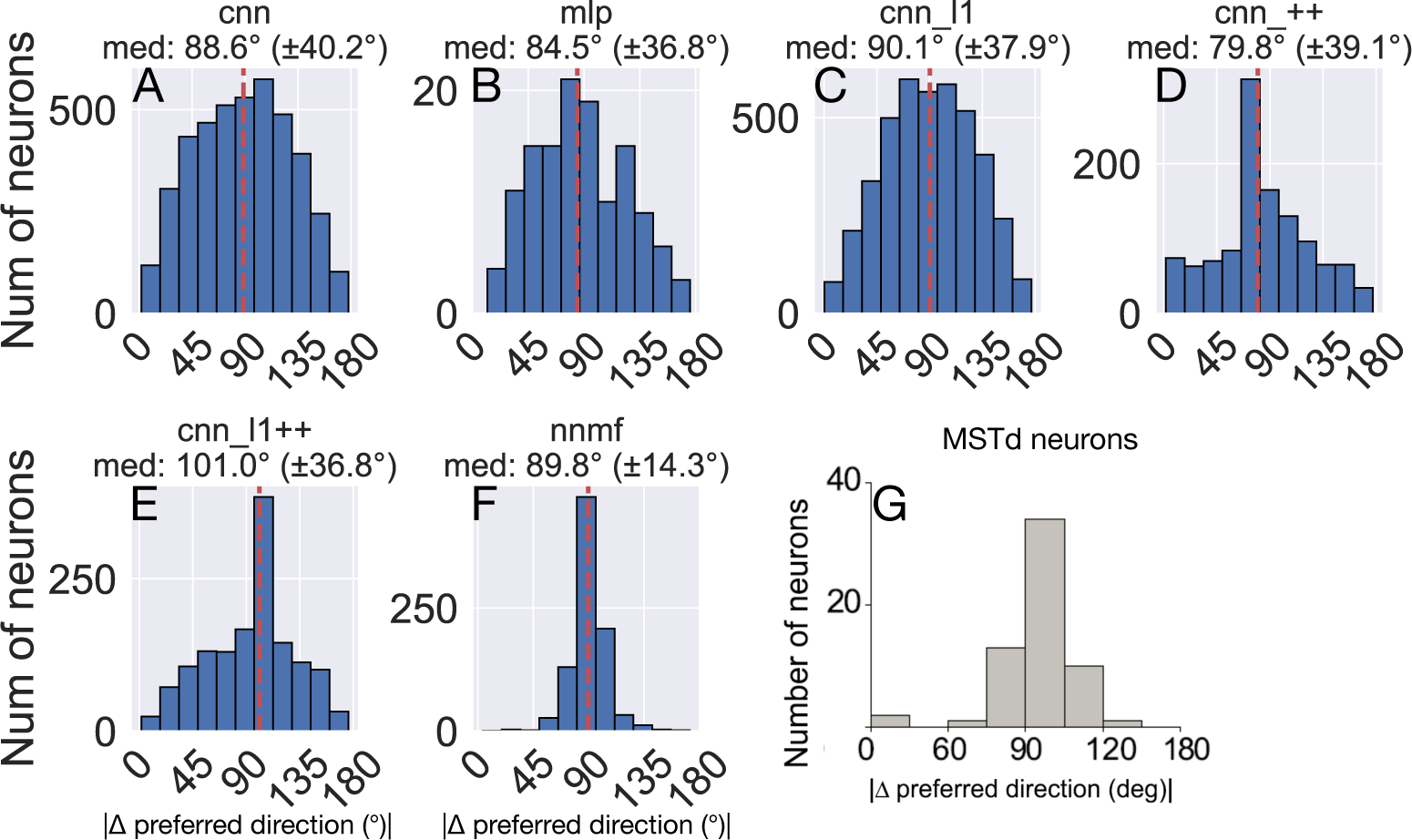
Difference between each neuron’s translation and rotation preference. The values above each plot show the median difference across the population as well as its standard deviation. (G) Difference in translation and rotation preference in MSTd population from Takahashi et al. (2007). The dashed vertical line corresponds to the median difference in preference.

### Rotation tuning index

Figure 15 shows the rotation tuning index (RTI) of each model unit. The RTI measures the strength of a neuron’s rotation tuning on a scale between 0 and 1. We compute RTI in the same way as HTI (Eq. 9), except activations correspond to the TestProtocolR diagnostic dataset that contains rotation-only optic flow samples and the labels in Eq. 9 now correspond to rotation. All the models except for MLP possess similar median RTI values across the population of ≈0.25. This indicates weak overall rotation tuning. The MLP has a significantly larger median RTI, which appears to arise due to a sizable subpopulation with RTIs between 0.9–1.0 that selectively respond to a small number of rotation optic flow patterns (95% CI: MLP [0.61, 0.65] vs. CNN [0.24, 0.26], CNN L1 [0.19, 0.22], CNN ++ [0.24, 0.26], CNN L1++ [0.26, 0.29], NNMF [0.25, 0.26]).

**Figure 15:**
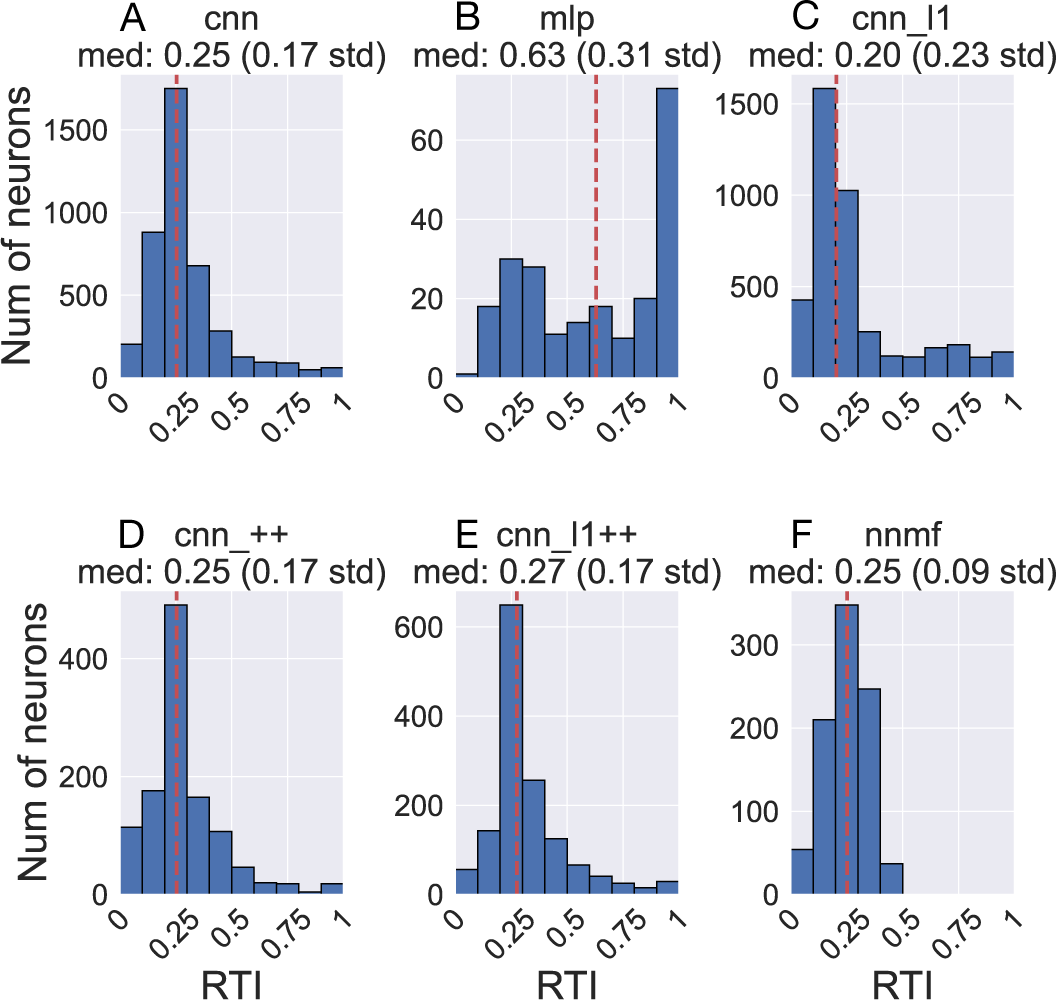
Histograms (10 bins) showing the rotation tuning index (RTI) of units in each model. Same format as Figure 11.

### Sparseness

Beyeler et al. (2019) propose that sparseness may represent a defining property of optic flow encoding in area MSTd. Following Beyeler et al. (2016), we computed sparseness metrics based on the model activations to optic flow samples in the TR360 test set (Vinje & Gallant, 2000). The population sparseness measures the proportion of neurons that activate to a single stimulus. The lifetime sparseness measures the proportion of stimuli in the dataset to which a single neuron activates. Both metrics range from 0 to 1. Zero indicates a dense code wherein a stimulus activates every neuron, while one indicates a grandmother cell localist code wherein a stimulus activates only a single neuron. Hence, an intermediate value indicates a sparse code wherein a subset of neurons activate to a stimulus. Figure 16A shows that all models garner comparable population and lifetime sparseness metrics on optic flow in the TR360 test set. Interestingly, all the CNNs yield average sparseness metrics of ≈0.5, which is compatible with a sparse code. The NNMF average sparseness metrics indicate a denser code, while those garnered by the MLP indicate a more localist code.

**Figure 16:**
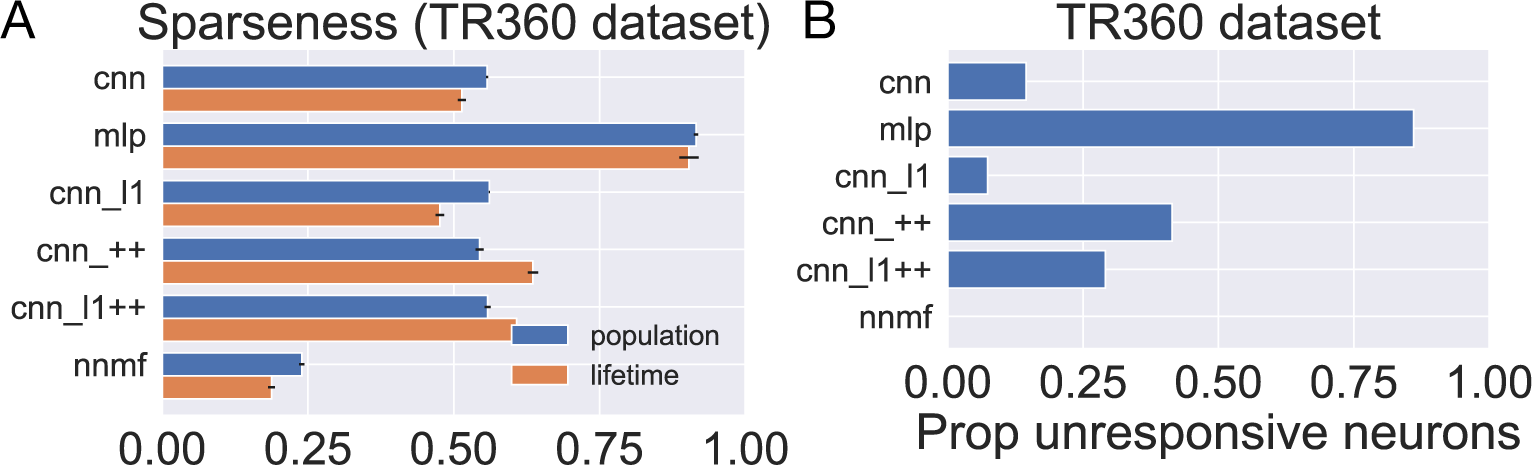
Measures of sparseness in the computational models computed on the TR360 test set. (A) Population and lifetime sparseness metrics. Error bars show 95% confidence intervals on the mean sparseness across neurons (population) or stimuli (lifetime) in each model. (B) The proportion of neurons that did not elicit responses to any of the stimuli in the TR360 test set.

We examined another phenomenon that promotes sparseness in deep neural networks that use the common ReLU activation function as we do here. Because neurons that implement the ReLU activation function output zero activation when their input is subthreshold, it is possible throughout the course of training for a neuron’s afferent weights to be set so that even the largest possible inputs cannot activate the neuron. This is known as the “dying ReLU” (Lu et al., 2019). While this is often referred to as a “problem” in the machine learning literature, this phenomenon may confer benefits, such as forcing the neural network to encode stimuli in a subset of available neurons. This may encourage sparseness, simplicity, and efficiency in the stimulus encoding (Glorot et al., 2011). Figure 16B shows the proportion of unresponsive neurons in each model, which we define as the proportion of neurons that never activate to any optic flow stimulus in the TR360 test set. There are 15% unresponsive neurons in the CNN. The CNN L1 model possesses half the percentage of unresponsive neurons (7%) and the CNN ++ and CNN L1++ models possess about double the percentage, 41% and 29%, respectively. The percentage is considerably greater in the MLP model (86%), which suggests a sparse encoding of optic flow. Differences in the number of neurons are unlikely to explain the disparity in unresponsive neurons since, for example, MLP, CNN ++, and CNN L1++ have roughly the same number — 2448, 2244, and 2291, respectively. Interestingly, all the neurons in the NNMF responded to at least one stimulus — it contains no unresponsive neurons. This likely stems from the computational goal of NNMF: to learn a small set of basis vectors that may be used to reconstruct the stimulus dataset.

Taken together, the sparseness metrics suggest that the CNNs learned a sparse optic flow encoding, the MLP learned an even sparser representation, and NNMF learned a denser code.

## Discussion

We investigated whether neurons in CNNs optimized to accurately estimate self-motion effectively model optic flow tuning properties in primate MSTd. In general, we find that such CNNs do not capture MSTd tuning properties as well as the NNMF algorithm that performs dimensionality reduction on neural motion signals. Our findings suggest that accurate self-motion estimation may not be a primary computational goal subserved by MSTd. This is surprising given that CNNs that are optimized to accurately classify natural images represent the leading models of neural primate ventral stream (Schrimpf et al., 2018; Yamins et al., 2014; Lindsay, 2021; Serre, 2019).

We observe an inverse relationship between accuracy of self-motion estimates and correspondence with MSTd-like tuning properties. The neural networks (CNN, MLP, CNN L1) that achieve the smallest mean errors of ≤10° and ≤1.5°/s when estimating translation and rotation, respectively, deviate substantially from MSTd translation preferences (Figures 8), peak heading discriminability (Figure 10), and rotation preferences (Figures 13). On the other hand, NNMF does not yield accurate self-motion estimates on the TR360 test dataset, yet it best captures the MSTd tuning properties in most analyses. Given that NNMF garners substantially improved self-motion decoding accuracy on the TR360 training set compared to the test set results shown in Figure 4, it is possible that the sparse representation may hinder generalization (Spanne & Jörntell, 2015). However, PCA does not produce a sparse representation and decoding from a PCA encoding of the test stimuli MT motion signals does not produce more accurate predictions. It is noteworthy that the sizeable disparity in accuracy between the CNNs and NNMF only emerges on the TR360 dataset (Figure 4) — the two model classes yield comparably good accuracy on the far simpler BenHamedT and BenHamedR datasets (Figure 5), which have been previously used to evaluate the decoding accuracy of NNMF (Beyeler et al., 2016). This raises the possibility that NNMF may struggle to encode the complexity of the TR360 with only 64 basis vectors. However, increasing the number of basis vectors to several hundred did not overcome this issue in our testing. The contrast in results between the TR360 and Ben Hamed datasets highlights the importance of validating models against benchmark tasks that capture a more naturalistic range of parameters. We are not aware of an existing neurophysiological study that reports the accuracy with which translation and rotation may be decoded from MSTd for optic flow stimuli as complex as those in the TR360 dataset. This would be valuable for testing the predictions of competing computational models.

What is the computational goal of MSTd if it is not accurate self-motion estimation? Beyeler et al. (2019) argue that nonnegativity and sparseness represent computational principles that give rise to a wide range of neural properties, including optic flow tuning in MSTd. Our findings suggest that these computational constraints may not be sufficient for capturing optic flow tuning in MSTd, at least while accurate self-motion estimation remains a primary goal. The inclusion of the sparseness constraint does not appear to greatly impact optic flow encoding relative to the baseline CNN across our analyses. For example, CNN L1, like the CNN model, yields substantial tuning for backward translation, which is absent in the MSTd data (Figure 8). This could be because the CNNs are not optimizing to achieve a certain level of sparseness. If the sparseness induced by greater L1 regularization inversely impacts the accuracy of self-motion estimation, then the hyperparameter search process should select networks with weak L1 regularization. Indeed, the relatively small, albeit nonzero, values in the optimized CNNs support this possibility (Table 3). Modifying the CNN cost function to prioritize sparseness over self-motion motion accuracy may yield more MSTd-like tuning properties (Kashyap, Fowlkes, & Krichmar, 2019; Layton & Fajen, 2022). Yet even when the hyperparameter search process selected networks with sparser representations, as is the case for MLP (Figure 16), the network does not capture MSTd properties any better than the CNN or CNN L1.

We do find, however, that the non-negativity weight constraint alone consistently yields more MSTd-like tuning, particularly in the case of translation. Nevertheless, the model translation and rotation tuning properties still deviate from the MSTd data in numerous meaningful ways. For example, the CNNs with the non-negativity constraint do not have nearly as many neurons that prefer rightward self-motion (Figure 8), are lacking in peak heading discriminability for backward translation (Figure 10), and far too many neurons have preferences close to the yaw rotation axis (Table 5). The CNN L1++ model that implements both sparseness and non-negativity constraints does not approximate the translation and rotation tuning characteristics of MSTd better than the CNN ++ model that incorporates only the non-negativity constraint. This indicates that the two factors do not appear to act synergistically in the CNNs simulated here.

One possible explanation for the lack of sufficiency of the sparseness and non-negativity constraints is that additional, yet-to-be-identified mechanisms are required to explain MSTd optic flow tuning. Indeed, NNMF yields the best overall agreement among the simulated models to MSTd properties, but some discrepancies remain (e.g. Figures 10, Figures 11). Maus and Layton (2022) found that accuracy-optimized CNNs do not match the accuracy of human heading perception from optic flow in important scenarios including in the presence of independently moving objects, scenes that produce sparse optic flow fields, and in the presence of visual rotation. Processing optic flow over time with recurrent connections improved the consistency with human self-motion judgments, so the inclusion of this mechanism might also yield more MSTd-like tuning properties. It is noteworthy, however, that NNMF achieves much better consistency with MSTd than the CNNs without processing optic flow over time.

Alternatively, the discrepancy between the CNNs and MSTd may stem from assumptions within the selected deep learning framework. Training CNNs on a dataset with a large number of labels using a supervised learning paradigm may align well with the computational objectives of ventral stream (Khaligh-Razavi & Kriegeskorte, 2014), but perhaps not those of the dorsal stream. For example, it has been argued that the self-supervised learning paradigm represents a more biologically plausible learning method since it relies on stimulus-derived predictions to drive learning in lieu of ground truth labels (Halvagal & Zenke, 2023; Rao & Ballard, 1999; Mineault, Bakhtiari, Richards, & Pack, 2021). Another possibility is that supervised learning based neural networks may be compatible with the dorsal stream, but the goal is something other than achieving accurate self-motion estimation, as we assumed here. For example, MSTd may participate in a larger system that optimizes perception-action objectives that subserve successful dynamic interactions with the environment, such as navigation toward goals (Page, Sato, Froehler, Vaughn, & Duffy, 2015; Page & Duffy, 2018; Alefantis et al., 2022).

## Conclusion

In conclusion, the modeling approach which best captures MSTd response properties (NNMF) is unable to accurately decode heading and rotation from a more naturalistic and complicated dataset (TR360) than those previously considered (BenHamedT, BenHamedR). When examining accuracy-optimized neural networks, which performed better on TR360, we find an inverse relationship between accuracy of self-motion estimates and correspondence with MSTd-like tuning properties. Adding a nonnegativity constraint to the CNNs induces more MSTd-like sensitivity while adding L1 regularization had little effect. Our findings suggest that (1) some computational constraints are unaccounted for, such as a minimum amount of L1 regularization or another form of sparsity, or (2) the cost function for the accuracy-optimized networks is unaligned with the computational goals of primate MSTd.

## Acknowledgements

This article has been authored by an employee of National Technology & Engineering Solutions of Sandia, LLC under Contract No. DE-NA0003525 with the U.S. Department of Energy (DOE). The employee owns all right, title and interest in and to the article and is solely responsible for its contents. The United States Government retains and the publisher, by accepting the article for publication, acknowledges that the United States Government retains a non-exclusive, paid-up, irrevocable, world-wide license to publish or reproduce the published form of this article or allow others to do so, for United States Government purposes. The DOE will provide public access to these results of federally sponsored research in accordance with the DOE Public Access Plan https://www.energy.gov/downloads/doe-public-access-plan.

This paper describes objective technical results and analysis. Any subjective views or opinions that might be expressed in the paper do not necessarily represent the views of the U.S. Department of Energy or the United States Government.

